# A computational SOX10 network-based selection strategy to identify new drug targets in uveal melanoma

**DOI:** 10.1101/2025.10.14.679939

**Authors:** Anja Wessely, Christopher Lischer, Adrian Weich, Claudia Kammerbauer, Esther Güse, Elias A.T. Koch, Xin Lai, Jan Dörrie, Michael Erdmann, Caroline Voskens, Stefan Schliep, Jens Neumann, Markus Eckstein, Harald Knorr, Beatrice Schuler-Thurner, Julio Vera, Carola Berking, Markus V. Heppt

## Abstract

Uveal melanoma (UM) is the most common intraocular malignancy in adults. In contrast to cutaneous melanoma (CM), effective treatment options for metastatic UM are limited. The transcription factor SOX10 is crucial for CM initiation and survival, making it an interesting candidate for new targeted therapies, but its relevance in UM was unclear. We found that SOX10 was widely expressed in UM and essential for proliferation, cell cycle progression, and survival. The effects were partially mediated by SOX10-related genes including *MITF*, highlighting high addiction of UM to the SOX10-MITF axis. Additionally, SOX10 knockdown induced massive transcriptomic changes. Due to a lack of specific inhibitors of SOX10 and MITF, a computational approach was used to identify druggable targets by curating a UM-specific protein interaction network to search for candidates downregulated upon SOX10 inhibition. Thereby, the E2F transcription factor family was identified and their potential as druggable target candidates in UM was confirmed using the pan-E2F inhibitor HLM006474, resulting in cell cycle arrest and apoptosis. Taken together, SOX10 is crucial for UM survival and SOX10-associated proteins may serve as promising targets for developing new therapeutic strategies in UM.

## Introduction

Uveal melanoma (UM) is the most common intraocular tumor in adults and arises from melanocytes residing in the choroid, ciliary body, or iris ^1^. However, UM is by far less common than cutaneous melanoma (CM) and regarded as an orphan cancer ^2^. Although melanocytes represent the cells of origin of both UM and CM, these melanoma subtypes significantly differ regarding their tumor biology, clinical features, and therapy options ^3^. Despite successful local tumor control that can be achieved by radiotherapy or enucleation, almost 50% of patients eventually develop metastases for which successful treatments are sparse ^4,5^. In contrast to CM, immune checkpoint blockade (ICB) using anti-CTLA-4 and anti-PD-1 antibodies achieved low response rates in metastatic UM, presumably due to its low tumor mutational burden ^6,7^. UM lacks mutations in *BRAF* and *NRAS*, which are common in CM. Thus, small molecule inhibitors such as BRAF inhibitors that are successfully used in CM are ineffective in UM ^3^. Instead, more than 80% of UM carry mutations in genes coding for small G proteins GNAQ and GNA11, resulting in constitutive activation of MAPK signaling ^8,9^. However, targeted therapies inhibiting for instance MEK activity did not lead to any survival benefits in metastatic UM ^10,11^. As UM predominantly metastasizes to the liver, several liver-directed therapies have been developed which can prolong the survival of the patients, but the prognosis of these patients is still poor and most of them die within one year after diagnosis of metastases ^12^. So far, the bispecific molecule tebentafusp is the only systemic therapy that has demonstrated a benefit regarding overall survival, but can only be used in the subset of HLA-A*02:01-positive patients due to its molecular design ^13^. Thus, there is a high unmet medical need for developing new treatment options for metastatic UM patients.

Melanocytes are the pigmented cells of the body that are mainly present in the epidermis in the skin but also in other organs such as the uveal tract of the eye, brain, inner ear, and even in the heart ^14^. They develop from melanocytic precursor cells derived from the neural crest (NC), a highly motile transient cell population that forms during neurulation between the ectoderm and the neural tube within the first weeks of embryonic development ^15,16^. From the NC, melanocytic precursor cells termed melanoblasts migrate to their final destinations, e.g., the epidermis where they develop into mature melanocytes ^16^. Delamination, migration, cell survival, and maturation of these cells are tightly controlled by various NC-related transcription factors including sex-determining region Y-bo× 10 (SOX10) and MITF ^17^. The expression of transcription factors that are important during embryonic development can be maintained or reinduced in tumors to promote tumor initiation and progression, as e.g., Brn3a and MSX1 in CM ^18, 20^. SOX10 expression is essential for cell survival during melanoblast migration, but it is maintained in adult mature melanocytes, benign nevi, and also CM ^21, 26^. SOX10 is involved in CM formation and its inhibition reduced cell viability, induced cell cycle arrest, and apoptosis or senescence ^24, 26^. Thus, SOX10 could be an attractive candidate for the development of new targeted therapy approaches in melanoma. SOX10 expression was also found in UM where it may serve as a sensitive marker to distinguish UM from adenocarcinomas or adenomas derived from pigmented ciliary epithelium (APCE) ^27,28^ and one case of a UM patient harboring a 20 base pairs (bp)-spanning deletion in the *SOX10* gene was reported ^29^. Apart from this, SOX10 has been hardly investigated in UM and especially its functional role in UM remained unclear so far.

In this study, we show that SOX10 is widely expressed in patient-derived specimens and cell lines and crucial for UM cell survival. We identified its target gene *MITF* as one of the major mediators of the pro-survival effects. To translate these results into novel therapeutic approaches, we used an innovative computational network-based target selection strategy to identify druggable proteins which can easily be inhibited by small molecule inhibitors instead of SOX10 or MITF. Following this approach, the E2F transcription factor family was identified and confirmed as suitable target candidates in proof-of-concept *in vitro* experiments.

## Methods

### Analysis of SOX10 expression in UM tumor samples

Gene expression of *SOX10, POU4F1, MSX1*, and *SOX9* was explored by analyzing RNA sequencing data of 80 UM primary tumors from the TCGA *Ocular Melanomas* dataset ^30^ (accessed via UCSC Xena Browser ^31^) and of tumor samples obtained from two independent patient cohorts. The tumor samples of the first cohort (UM primary tumors: n=12, liver metastases: n=3) of UM patients from the Uniklinikum Erlangen were analyzed using the Nanopore^®^ sequencing technology. The tumor samples of the second independent cohort included 16 metastases (liver: n=14, breast: n=1, lung: n=1) from UM patients who participated in a phase 1 clinical trial at the Uniklinikum Erlangen (NCT04335890) ^32^. Gene expression data were visualized in heat maps created with the web application Heatmapper ^33^.

Immunohistochemical staining was used to explore SOX10 protein expression in 42 formalin-fixed and paraffin-embedded (FFPE) tumor samples (liver: n=28, soft tissue: n=5, orbita: n=2; greater omentum, ear, breast, lymph node, adrenal glands, kidney, lower back: n=1 each) obtained from the Department of Pathology, Uniklinikum Erlangen, and the Department of Pathology, Ludwig-Maximilian University Munich (both Germany). Detailed information about RNA sequencing and immunohistochemical staining procedures are described in the Additional file A1.

### Cell lines

UM cell lines 92.1 (RRID:CVCL_8607), Mel270 (RRID:CVCL_C302), OMM1.5 (RRID:CVCL_C307), and Mel285 (RRID:CVCL_C303) were kind gifts from Prof. Klaus Griewank, University Hospital Essen, Germany. UM cell lines OMM-1 (RRID:CVCL_6939), OMM2.3 (RRID:CVCL_C306), OMM2.5 (RRID:CVCL_C307), Mel202 (RRID:CVCL_C301), and Mel290 (RRID:CVCL_C304) were kindly provided by Martine Jager, University of Leiden, The Netherlands, and CM cell line 1205Lu (RRID:CVCL_5239) was a kind gift from Meenhard Herlyn, The Wistar Institute, Philadelphia, USA. Authentication of all UM cell lines and CM cell line 1205Lu was performed by a commercial provider (Eurofins Genomics, Ebersberg, Germany). Details about cultivation are presented in the Additional file A1.

### siRNA transfection for knockdown experiments

Cells were seeded in 6-well plates in the respective media lacking 1x antibiotic-antimycotic supplement and transfected on the following day with 20 nM of SOX10- or MITF-specific siRNAs or control siRNA (Supplementary Table 1) using 1.25 µl Lipofectamine RNAiMAX (Invitrogen, for UM and CM cells) per well or jetPRIME® DNA & siRNA Transfection Reagent (Polyplus-transfection S.A., Illkirch, France, for HM transfection) following the manufacturer’s instructions. All cells were cultivated at 37°C and 5% CO_2_ atmosphere in a humidified incubator for 24 h up to 96 h.

### Assessment of cell viability and cell death

Cell viability was assessed in cells grown in 6-well plates (knockdown experiments) or 24-well plates (inhibitor treatment) after 24 h up to 96 h of incubation using the CellTiter-Blue® Cell Viability Assay (Promega, Madison, Wisconsin, USA) according to the manufacturer’s protocol. The fluorescence intensity of the supernatant (excitation: 530 nm, emission: 590 nm) was determined in a plate reader (CytoFluor™ 2350 (Millipore) and Glo-Max® Explorer (Promega) for knockdown experiments; Infinite 200 PRO (Tecan, Grödig, Austria) for inhibitor experiments), and relative cell viability compared to respective control cells was calculated in Microsoft Excel. For apoptosis analysis, cells from knockdown or inhibitor experiments were stained with FITC-labelled annexin V using the Annexin-V-FLUOS Staining Kit (Roche, Penzberg, Germany) according to the manufacturer’s protocol and 0.5 µg/ml propidium iodide (Sigma-Aldrich, Taufkirchen, Germany). Flow cytometry analyses were performed after 48 h (knockdown experiments) on a FACScan cytometer (Becton Dickinson, Franklin Lakes, New Jersey, USA) and analyzed using the software CellQuest (Becton-Dickinson, Heidelberg, Germany) or on a FACSCanto II Cytometer after 72 h (inhibitor experiments) and analyzed using the software FlowJo, version 10.8.1 (both from BD Biosciences, San Jose, California, USA).

### Cell cycle analysis

Cells were seeded in 6-well plates in the respective media lacking 1x antibiotic-antimycotic supplement and transfected on the following day as described above. For inhibitor experiments, 50 µM HLM006474 (Selleck Chemicals LLC, Houston, Texas, USA) or equal amounts of DMSO (Sigma-Aldrich) as control were added the day after seeding. Cell cycle analysis was carried out as described previously by Graf et al. ^26^ after 48 h (knockdown experiments) on a FACScan cytometer and analyzed using the software CellQuest or after 72 h (inhibitor experiments) on a FACSCanto II Cytometer and analyzed using FlowJo, version 10.8.

### Analysis of transcriptomic changes in SOX10 knockdown cells

To determine the transcriptomic changes after SOX10 inhibition, UM cell lines 92.1 and Mel270 were seeded in T75 cell culture flasks one day prior to transfection with siSOX10-B or control siRNA. Total RNA was extracted 24 h after transfection and SOX10 knockdown efficacy was confirmed by qPCR analysis. RNA sequencing analysis of pooled total RNA derived from four individual experiments was performed by a commercial provider (CeGaT, Tübingen, Germany). Further information about RNA isolation and sequencing is provided in the Additional file A1.

### Reconstruction of a SOX10-based UM-specific protein interaction network

The most important intracellular signaling pathways in UM were identified in a literature survey (Supplementary Table 2) and the corresponding components were downloaded from the Reactome database and imported to CellDesigner (version 4.4.2). All entities were annotated directly with the Minimal Information Required In the Annotation of Models (MIRIAM) interface ^34^. Moreover, MITF, DCT, TYR, KIT, IRF4, MEF2C, ERBB3, ITPR2, CEBP, CREB3L2, BHLHB2, ETS1, E2F1, TYRP1, NES, EDNRB, PAX3, RET, EGR2, POU3F1, POU3F2, SP1 and MED1 which have been previously described to interact with SOX10 ^35,36^, were added. Proteins in the network were annotated with their UniProt ID, genes with their Ensembl ID, and miRNA with their miRBase ID. The manually curated network was then automatically extended with database knowledge using miRNexpander and miRTARBase (version 6.1), miRecords (version 4.5), HTRIdb (version 1), and TRANSFAC (version 2015.1), and species restrictions were set to human. The expanded network was further processed using Cytoscape (version 3.8.0). To include even more SOX10-interacting proteins and interactions among them, the expanded UM network was merged with our previously established SOX10 interaction network based on the oligodendrocyte differentiation cascade ^37^, duplicated nodes and self-loops were then removed and the network was pruned with published expression data from 63 UM microarrays (GEO ID: GSE22138) ^38^ and RNA sequencing data of 80 UM primary tumors (TCGA *Ocular Melanomas* dataset ^30^) as a second dataset. The expression values were converted to transcripts per million (TPM), a mean value for each gene was calculated, mapped separately to the network nodes via their genes Ensembl IDs, and all nodes with an average expression value greater than 1 in either dataset were selected. Afterwards, network analysis and topology statistics were computed using the Network Analyzer in Cytoscape. For the extraction of the regulatory core of the network, the UM network was queried for regulatory motifs using the Cytoscape application NetMatchStar and seven types of these regulatory motifs were examined. To detect the most important nodes and their interactions, a weighted ranking score of the previously identified motifs was calculated (see Additional file A1 for details).

### Statistics

All experiments were performed in triplicates unless otherwise stated. The presented data are shown as mean ± standard deviation (SD). Student’s T-test was used for comparing two groups and one-way ANOVA with Dunnett’s Post Hoc-Test was used for multiple comparisons. Densitometric analyses of immunoblot images were performed with Image J (Fiji). The correlation between SOX10 and MITF protein expression was evaluated with Pearson’s correlation. Calculations were performed in SPSS Statistics (version 28, IBM, Armonk, New York, USA) or GraphPad Prism (version 5, GraphPad Software, Inc., San Diego, California, USA). A p-value of < 0.05 was considered significant unless otherwise stated.

### Further methods

Further methods are described in Additional file A1.

## Results

### SOX10 is widely expressed in patient-derived primary tumors, metastases, and UM cell lines

NC-related transcription factors are important for CM development and progression, but their role in UM has not been investigated so far. Therefore, we analyzed the gene expression of *SOX10, SOX9, POU4F1* (coding for Brn3a), and *MSX1* in the publicly available *TCGA Ocular Melanomas* dataset containing transcriptomic data derived from 80 UM primary tumors ^30^. *SOX10* gene expression was markedly higher in all samples compared to other NC-related transcription factors such as *POU4F1* and *MSX1* or the closely related transcription factor *SOX9* (Figure 1A). These results were confirmed when analyzing RNA sequencing data of two independent UM patient cohorts from the Uniklinikum Erlangen (Figure 1B and C). Similar to the gene expression analyses, we observed strong nuclear SOX10 staining in the analyzed samples obtained from liver metastases (Figure 1D; Figure S1A, B) and distant metastases in other organs (Figure S1C, D, E). More precisely, we found > 1% SOX10 positive cells in 37 out of 42 analyzed samples (88.1%) and > 10% SOX10 positive cells in 26 out of 42 analyzed samples (61.9%) (Figure 1F, G), indicating that SOX10 protein expression is maintained in UM metastases at a high expression level. Next, we analyzed SOX10 gene and protein expression in UM cell lines derived from primary tumors and metastases, human melanocytes (HM), and the CM cell line 1205Lu that has already been previously analyzed by our group ^26,39^. High SOX10 expression was detected in seven out of nine analyzed UM cell lines and was higher compared to CM cell line 1205Lu (Figure 1E, F). However, no SOX10 expression was observed in UM cell lines Mel285 and Mel290, which also do not express melanocytic markers such as Melan A, gp100, TYR, TYRP1, and DCT ^40^. Taken together, these results indicate that SOX10 is widely expressed in both UM primary tumors and metastases, and SOX10 expression is maintained in most UM cell lines, suggesting that SOX10 may be a necessary factor for UM survival and progression.

**Figure 1:**
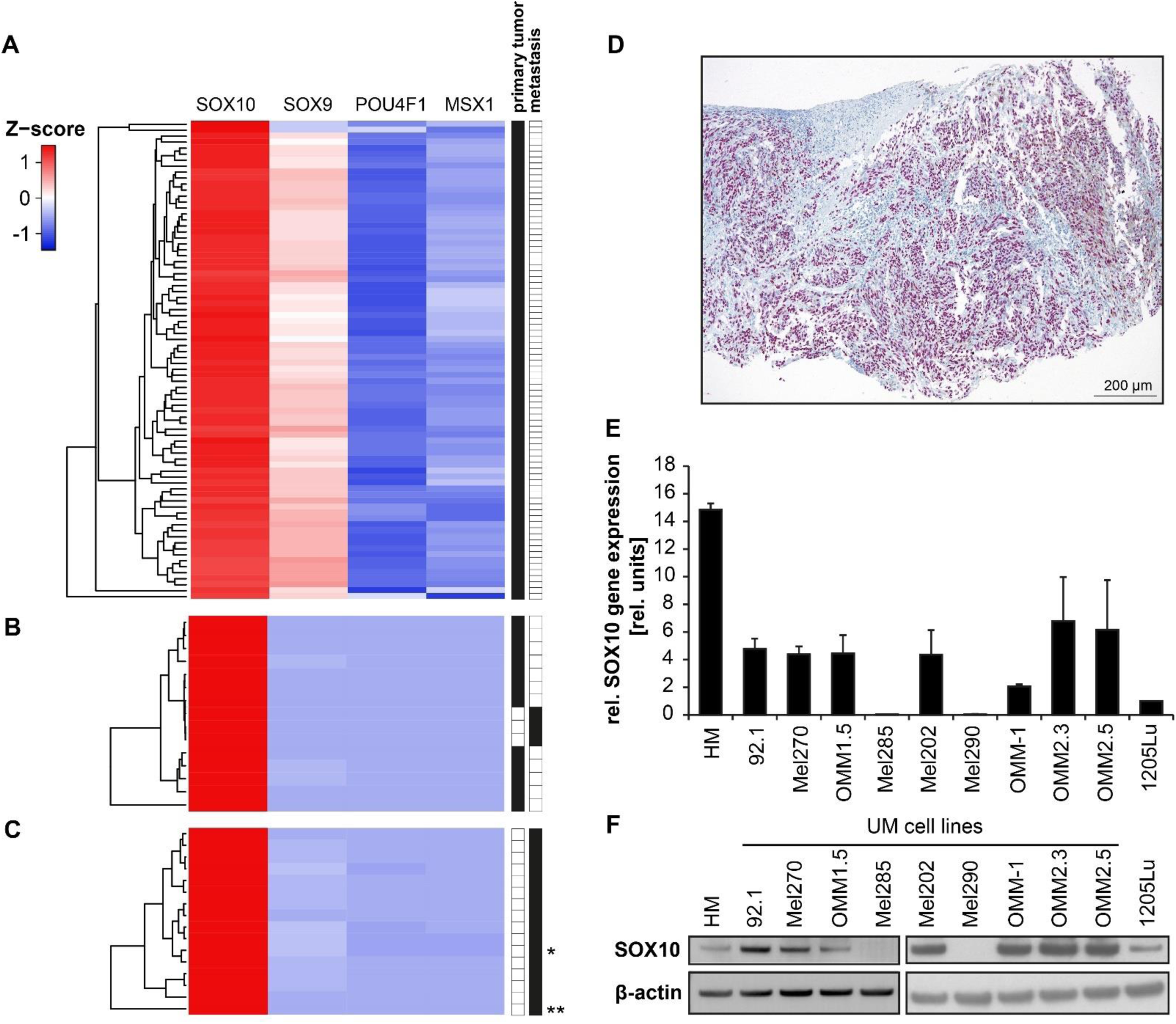
SOX10 is widely expressed in UM primary tumors, metastases, and cell lines. Heat maps showing gene expression of *SOX10, SOX9, POU4F1* (coding for Brn3a protein), and *MSX1*, data obtained from (A) the TCGA *Ocular Melanomas* dataset including RNA sequencing data of 80 UM primary tumors (accessed via UCSC Xena Browser) ^31^, (B) 12 UM primary tumors and three liver metastases obtained from a patient cohort of UM patients from the Uniklinikum Erlangen and (C) metastases derived from the liver (n=14), lung (*, n=1), and breast (**, n=1) of UM patients participating in a phase 1 clinical trial investigating IKKb-matured, RNA-loaded dendritic cells for the treatment of metastasized UM. (D) Representative microscopic image of immunohistochemical SOX10 staining of a UM liver metastasis, scale bar = 200 µm. (E) Relative SOX10 mRNA expression levels in human melanocytes (HM), UM cell lines 92.1, Mel270, OMM1.5, Mel285, Mel202, Mel290, OMM-1, OMM2.3, OMM2.5, and CM cell line 1205Lu, normalized to GAPDH expression. The relative expression level of 1205Lu was set to 1, data represent mean ± SD, n=3. (F) SOX10 protein expression in human melanocytes (HM), UM cell lines 92.1, Mel270, OMM1.5, Mel285, Mel202, Mel290, OMM-1, OMM2.3, OMM2.5, and CM cell line 1205Lu. β-actin served as loading control.

### SOX10 inhibition decreases cell viability and leads to cell cycle arrest and apoptosis

To characterize its functional role in UM, SOX10 expression was inhibited via RNA interference in the three SOX10 high-expressing UM cell lines 92.1, Mel270 (both derived from primary tumors), and OMM1.5 (derived from a liver metastasis of the same patient as Mel270, equal to OMM2.5 ^41^) as well as in HM and CM cell line 1205Lu. We observed a massively decreased cell viability in UM cell lines 92.1 and Mel270 already 48 h after siSOX10-A or siSOX10-B transfection compared to controls and a significant decrease in OMM1.5 similar to CM cell line 1205Lu after 72 h and 96 h (Figure 2A). Interestingly, SOX10 inhibition did not affect HM cell viability. Consistent with this, SOX10 knockdown led to changes in cell morphology from a flat, cobblestone-like, or star-shaped to a round phenotype, decreased cellular density, and cells started to detach from the cell culture dish surface, indicating that cell death occurred (Figure S2A). These alterations were not observed in HM. Propidium iodide staining revealed that SOX10 knockdown induced an arrest in the G1 phase of the cell cycle in UM cells and CM cell line 1205Lu, but not in HM (Figure 2B). Hypophosphorylation of the retinoblastoma protein (Rb) in all tested UM cell lines (Figure 2C) and an increase in p21 and p27 expression in some UM cell lines was observed in immunoblot analyses (Figure S2B), confirming the findings of the flow cytometry analyses. To determine which type of cell death had occurred 48 h after siSOX10 transfections, annexin V-propidium iodide staining was performed. Loss of SOX10 massively increased the number of annexin V-positive cells in UM cell lines 92.1 and Mel270, indicating that SOX10 knockdown triggered apoptosis in these cell lines (Figure 2D). In contrast, SOX10 knockdown did not lead to an increase in apoptosis in UM cell line OMM1.5 as well as HM and CM cell line 1205Lu, which may be explained by the fact that the decrease in cell viability was not as pronounced as in 92.1 and Mel270 at that time point. Immunoblot analyses of UM cells 48 h after siSOX10 transfection revealed cleavage of caspase 9 in all cell lines and caspase 3 cleavage in 92.1 and Mel270 (Figure 2E), confirming the flow cytometry results. We also observed increased Ser139 phosphorylation of histone variant H2A.X (γ-H2A.X), a marker for DNA double-strand breaks, in 92.1 and Mel270 in SOX10 knockdown cells, suggesting that SOX10 inhibition may cause DNA damage. However, time course analyses showed that the increase in γ-H2A.X levels occurred later than the cleavage of caspase 3 and PARP (Figure S2C), indicating that apoptosis-related DNA fragmentation may have triggered the formation of γ-H2A.X as a secondary effect. Blocking of caspase 3 activity by the irreversible inhibitor Z-DEVD-FMK increased the cell viability of siSOX10-transfected cells and decreased caspase 3-mediated cleavage of PARP, which is typical for apoptosis (Figure S2D, E). Interestingly, p53 expression did not change upon SOX10 inhibition, indicating that cell cycle arrest and apoptosis are mediated in a p53-independent manner (Figure S2F). The expression analysis of several pro- and antiapoptotic proteins did not reveal clear differences between SOX10 knockdown and control cells (Figure S2F), underlining the complexity of SOX10’s role in cell death initiation. ERK, p38, and Akt signaling contribute to proliferation, survival, and mediate stress responses in cells ^42, 44^, and mutated *GNAQ* or *GNA11*, which are often found in UM, lead to constitutive activation of MAPK signaling ^8,9^. Thus, we explored if the loss of SOX10 expression alters the activation levels of these signaling pathways. SOX10 inhibition induced p38 stress kinase signaling in all tested UM cells and diminished Akt signaling whereas ERK activity levels were altered in a cell line-dependent manner (Figure S2G). Taken together, SOX10 inhibition decreased cell viability and cell proliferation, promoted cell cycle arrest in the G1 phase, activated p38 stress kinase signaling, diminished Akt signaling, and led to cell death via apoptosis in UM cells, indicating that SOX10 is crucial for UM cell survival.

**Figure 2:**
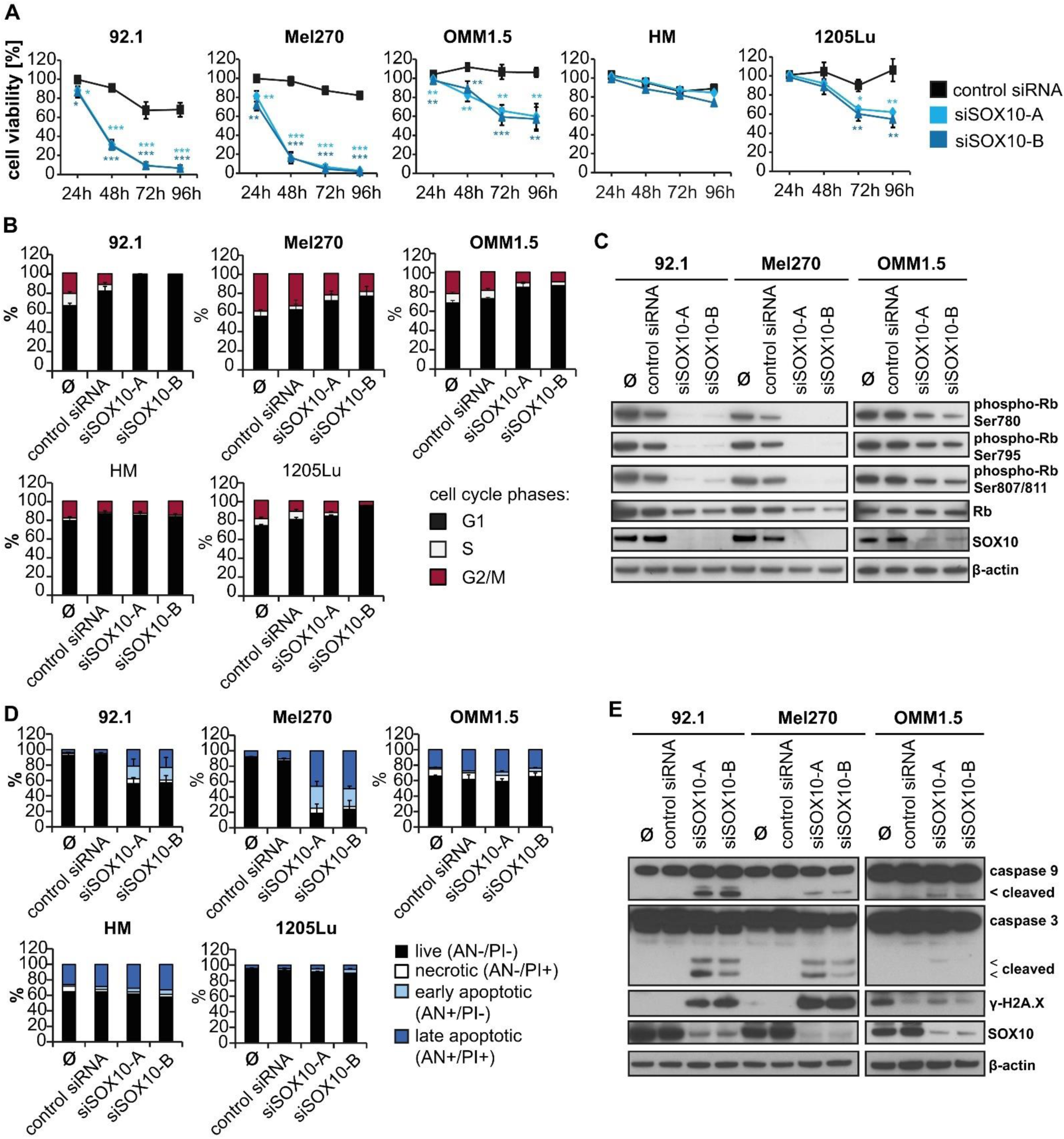
SOX10 inhibition decreases cell viability, leads to cell cycle arrest, and triggers cell death. (A) Cell viability of HM, 92.1, Mel270, OMM1.5, and 1205Lu transfected with control siRNA, siSOX10-A, or siSOX10-B for 24 h, 48 h, 72 h, and 96 h, mean ± SD, n = 3. *: p < 0.05, **: p < 0.01, ***: p < 0.001 vs. control siRNA. (B) Cell cycle analysis in HM, 92.1, Mel270, OMM1.5, and 1205Lu transfected with control siRNA, siSOX10-A, or siSOX10-B for 48 h. Mean ± SD, n = 3. *: p < 0.05, **: p < 0.01, ***: p < 0.001 vs. control siRNA. (C) Protein expression analysis of retinoblastoma protein (Rb) phosphorylation (phospho-Rb) at serin residues Ser780, Ser795, Ser807/811, total Rb, and SOX10 in UM cell lines 24 h after transfection with control siRNA, siSOX10-A, or siSOX10-B. (D) Apoptosis analysis by flow cytometry in HM, 92.1, Mel270, OMM1.5, and 1205Lu transfected with control siRNA, siSOX10-A, or siSOX10-B for 48 h. AN: Annexin V, PI: propidium iodide. Mean ± SD, n = 3. **: p < 0.01, ***: p < 0.001 vs. control siRNA. (E) Protein expression of caspases 9 and 3 (including cleaved fragments), histone variant H2A.X phosphorylated at Serin 139 (γ-H2A.X), and SOX10 in 92.1, Mel270, and OMM1.5 48 h after transfection with control siRNA, siSOX10-A, or siSOX10-B. β-actin served as loading control for immunoblots.

### SOX10 target gene MITF contributes to pro-survival effects of SOX10

In order to further decipher the mode of action by which SOX10 mediates its pro-survival functions, we focused on the transcription factor MITF, as its transcription is directly regulated by SOX10 ^45,46^. As “master regulator of melanogenesis”, MITF controls the transcription of genes that are involved in the melanin synthesis cascade including *TYR* and *DCT*, and has been described to mediate antiapoptotic functions ^47^. MITF was highly expressed in all tested UM cells except for Mel285 and Mel290 (Figure 3A, B), and SOX10 and MITF protein expression levels were significantly correlated (Pearson correlation coefficient r = 0.792, p = 0.004) (Figure S3A). As expected, SOX10 knockdown significantly reduced MITF gene and protein expression (Figure 3C; Figure S3B, C). To explore if MITF knockdown may have a similar pro-apoptotic effect as SOX10 knockdown, we transfected UM cells with siMITF and observed a decrease in UM cell viability (Figure 3D, Figure S3D) which occurred slightly delayed in OMM1.5. In MITF low-expressing 1205Lu cells, MITF knockdown did not affect the cell viability, as expected. We also found an arrest in the G1 phase of the cell cycle (Figure 3E), which was confirmed by Rb hypophosphorylation, decreased cyclin D1 expression, and increased p21 expression (Figure 3F). Expression of p27 was hardly altered and expression of p53 did not change in any cell line, indicating that the cell cycle arrest occurred in a p53-independent manner. Annexin V-propidium iodide staining showed an increase in annexin V-positive cells in 92.1 and Mel270 48 h after transfection with siMITF, but no differences in OMM1.5 and 1205Lu (Figure 3G), which is in line with the observations of the cell viability analyses. Immunoblot analyses revealed caspase 9 cleavage and increased γ-H2A.X expression in all tested UM cell lines and a decrease in Bcl-2 and cleavage of caspase 3 and PARP in UM cell lines 92.1 and Mel270 upon MITF inhibition (Figure 3F). Expression of other pro- and antiapoptotic markers such as Bcl-w, Bak, and Bax varied in a cell line-dependent manner (Figure S3E). Next, we investigated if ectopic MITF expression can counteract the pro-apoptotic effects of SOX10 inhibition and transfected Mel270 cells with a MITF expression vector leading to the expression of a FLAG® (DDK)- and Myc-tagged MITF protein followed by siSOX10 transfection on the following day (Figure S3F, G). Indeed, we observed increased cell viability (Figure 3H) and a decreased cleavage of caspase 9, caspase 3, and PARP in cells expressing ectopic MITF (Figure 3I), suggesting that MITF is at least partly able to counteract cell death induction triggered by the loss of SOX10 expression. Altogether, the data indicate that SOX10 mediates its pro-survival functions via MITF, and MITF seems essential for UM cell survival, similar to SOX10.

**Figure 3:**
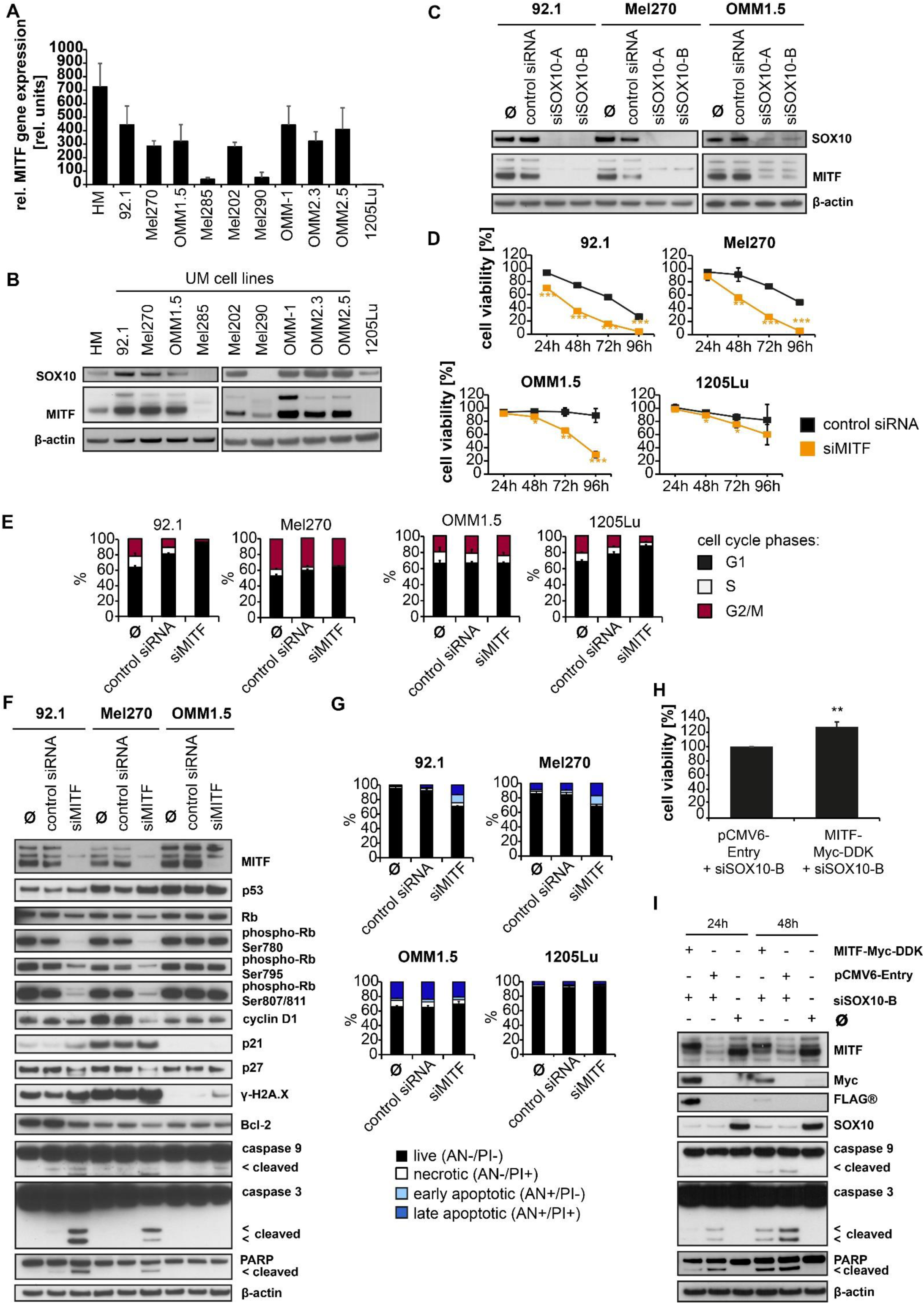
SOX10 target gene MITF contributes to pro-survival effects of SOX10. (A) Relative MITF mRNA expression levels in HM, 92.1, UM cell lines, and CM cell line 1205Lu, normalized to GAPDH expression. Relative expression level of 1205Lu was set to 1, mean ± SD, n=3. (B) Basal MITF and SOX10 protein expression in HM, UM cell lines, and CM cell line 1205Lu. (C) SOX10 and MITF protein expression 24 h after transfection with control siRNA, siSOX10-A, or siSOX10-B. (D) Cell viability after transfection with control siRNA or siMITF for 24 h, 48 h, 72 h, and 96 h, mean ± SD, n = 3. *: p < 0.05, **: p < 0.01, ***: p < 0.001 vs. control siRNA. (E) Cell cycle analysis after transfection with control siRNA or siMITF for 48 h. Mean ± SD, n = 3. (F) Protein expression of MITF, p53, total Rb, phosphorylated Rb (phospho-Rb) at Ser780, Ser795, Ser807/811, cyclin D1, p21, p27, histone variant H2A.X phosphorylated at Serin 139 (γ-H2A.X), Bcl-2, caspase 9 and caspase 3, and PARP (including cleaved fragments) after transfection with control siRNA or siMITF for 24 h. (G) Apoptosis analysis by flow cytometry after transfection with control siRNA or siMITF for 48 h. AN: Annexin V, PI: propidium iodide. Mean ± SD, n = 3. (H) Cell viability of UM cell line 92.1 transfected with empty control vector pCMV6-Entry or pCMV6-MITF-Myc-DDK expression plasmid followed by siSOX10-B transfection for 48 h, mean ± SD, n = 3. **: p < 0.01 vs. pCMV6-Entry + siSOX10-B. (I) Protein expression of MITF, Myc tag, FLAG® tag, SOX10, caspases 9 and 3, and PARP (including cleaved fragments) in Mel270 transfected with empty control vector pCMV6-Entry or pCMV6-MITF-Myc-DDK expression plasmid followed by siSOX10-B transfection for 24 h and 48 h. Ø: treatment with transfection reagents 3000 only. β-actin served as loading control for immunoblots.

### SOX10 knockdown massively alters the transcriptomic landscape in UM cells

As ectopic MITF expression did not completely antagonize the pro-apoptotic effects of the SOX10 loss, other proteins apart from MITF are likely involved in mediating the pro-survival functions of SOX10. Thus, we explored the overall changes in the transcriptome of SOX10-inhibited 92.1 and Mel270 cells by RNA sequencing analysis, revealing 6537 significantly up- and 5914 downregulated genes upon SOX10 inhibition (Figure 4A; Figure S4). We selected all genes with a |log_2_fold| differential gene expression of > 1 (1949 genes), and found that these genes are enriched in pathways related to cancer hallmarks such as epithelial-mesenchymal transition, hypoxia, angiogenesis, and apoptosis (Figure 4B). Similar pathways are identified for MITF target genes (Figure S5). Among the identified gene sets, 17 genes are predicted to be regulated by SOX10, and some of them are related to cell cycle and apoptosis. Notably, MITF was not among the genes on this list because it does not belong to any of the pathway terms shown in this figure. We further compared them with publicly available SOX10 chromatin immunoprecipitation (ChIP) seq data of melanocytes ^35^, and thereby identified 395 putative direct target genes of SOX10. GO enrichment analysis revealed that the direct SOX10 target genes are associated to pathways linked to developmental processes like “axonogenesis”, and “renal systems development”, but also to “peptidyl-tyrosine phosphorylation“/”peptidyl-tyrosine modification” (Figure 4C), and apoptosis (Figure 4D). Taken together, these analyses demonstrate that SOX10 inhibition massively alters the transcriptomic landscape in UM cells. Targets directly regulated by SOX10 were linked to developmental processes, and pathway enrichment considering all differentially expressed genes connected SOX10 knockdown to cell cycle arrest and apoptosis, suggesting that SOX10 may interfere or synergize with other central cancer transcription factors through a complex regulatory network.

**Figure 4:**
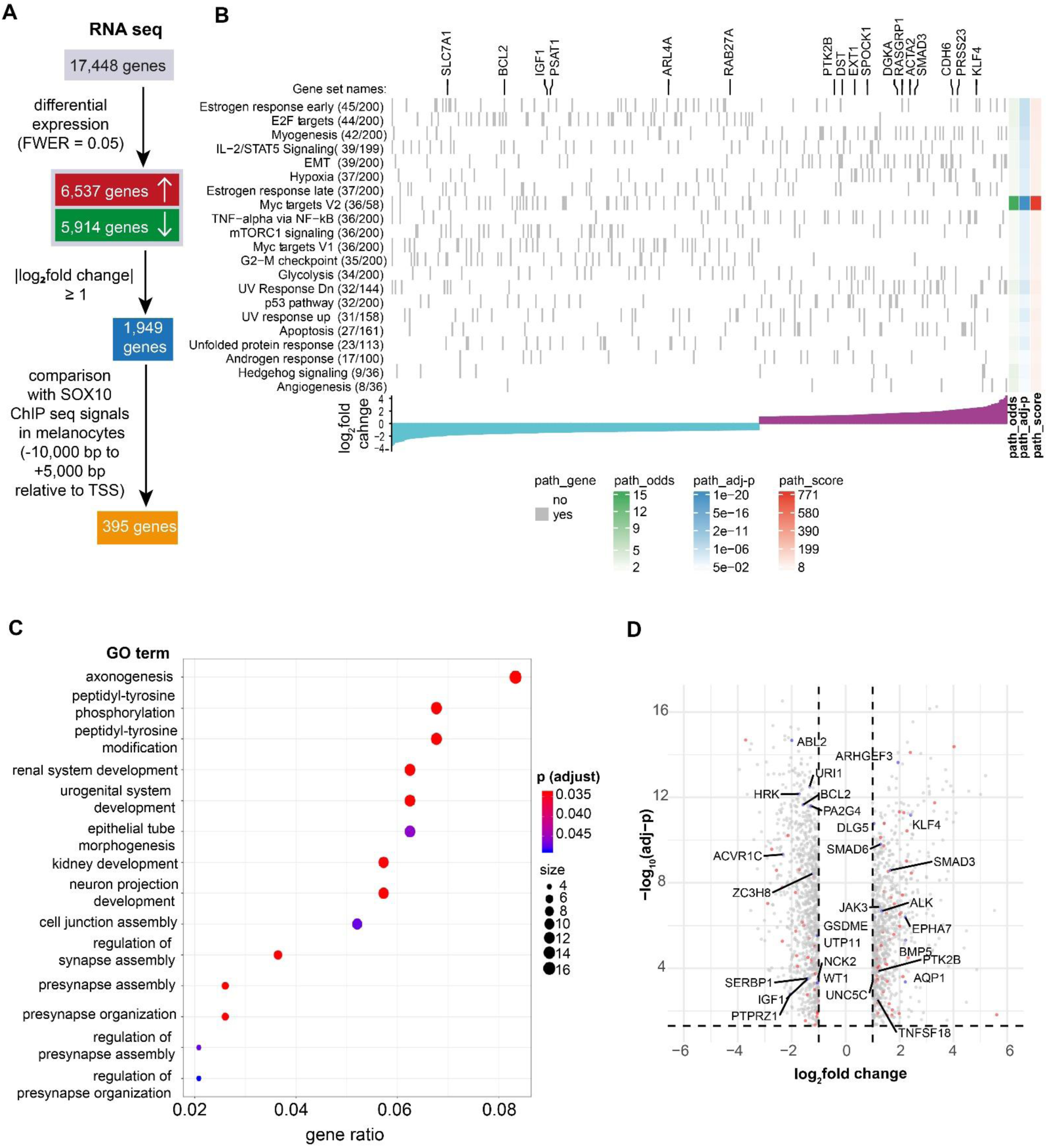
SOX10 knockdown massively alters transcriptomic landscape in UM cells. (A) RNA sequencing analysis workflow. UM cell lines 92.1 and Mel270 were transfected with control siRNA or siSOX10-B and total RNA was extracted after 24 h. After confirming SOX10 knockdown by qPCR analysis, the total RNA of four individual experiments was pooled and RNA sequencing performed, resulting in 17448 expressed genes. Overall, 5914 genes were significantly downregulated and 6537 significantly upregulated at a family-wise error rate (FWER) of 0.05. All genes with a |log_2_fold change| of ≥ 1 were compared to publicly available SOX10 ChIP seq data of melanocytes ^35^, identifying 395 putative direct target genes of SOX10. TSS: transcription start site. (B) Heat map showing the identified enriched hallmark gene sets from the MSigDB database. Only genes belonging to the identified enriched hallmarks are shown. The gene set names with the number of overlapping genes and the number of total genes in a gene set are shown on the left. Each gray grid in the heat map indicates whether a gene is enriched in a hallmark gene set. The odds ratio (odds), adjusted p-values (adjp) and a combine score (odds multiplied by [-log_10_(adjp)]) of each gene set are shown on the right. The bar graph at the bottom shows the log_2_fold change (cyan: downregulated; purple: upregulated) of the genes enriched in the gene sets. The 17 putative targets of SOX10 are shown at the top. (C) GO term analysis of the 395 putative direct target genes of SOX10 identified by RNA sequencing analysis after SOX10 knockdown. (D) Volcano plot showing the identified differentially expressed genes (|log_2_fold change| ≥ 1 and adj-p ≤ 0.05). Putative direct SOX10 target genes were highlighted in red and putative SOX10 target genes that are associated to apoptosis GO terms were labeled and highlighted in blue.

### Curating a SOX10-based UM-specific network for identifying candidates for targeted UM therapy

To date, the number of effective systemic treatments for metastatic UM patients is limited, and new therapy options are urgently needed. We showed that SOX10 is widely expressed in UM and crucial for the survival of UM cells but not for normal cells such as HM. Thus, the SOX10-MITF axis may be an ideal candidate for targeted therapy. However, SOX10 and MITF are mostly present in the nucleus ^48^, making it hardly accessible for blocking antibodies, and no specific inhibitors have been discovered yet. As shown above, inhibiting the SOX10 target gene *MITF* led to similar effects as direct siRNA-mediated SOX10 inhibition. Thus, we speculated that inhibiting druggable SOX10-associated proteins instead of directly inhibiting SOX10 or MITF themselves may exert similar pro-apoptotic effects. To search for such proteins, we applied a computational network-based and SOX10-centered target selection strategy. First, an UM core interaction network was created by combining manual curation of the literature and information from databases about proteins of important signaling pathways in UM, known SOX10-protein interactions, and miRNA-mRNA interactions to discover connections between relevant pathways in UM and SOX10-related proteins (Figure S6). We utilized a computational pipeline to expand this manually curated protein interaction network. Next, we applied a network biology algorithm to isolate a core regulatory network composed of highly interconnected and highly expressed molecular factors (65 nodes and 306 edges) (Figure S7). Finally, we expanded this core adding UM-relevant first neighbors of each node, reannotated, and filtered them according to their basic functions, leading to a new target selection network consisting of 394 nodes and 1620 edges (Figure 5).

**Figure 5:**
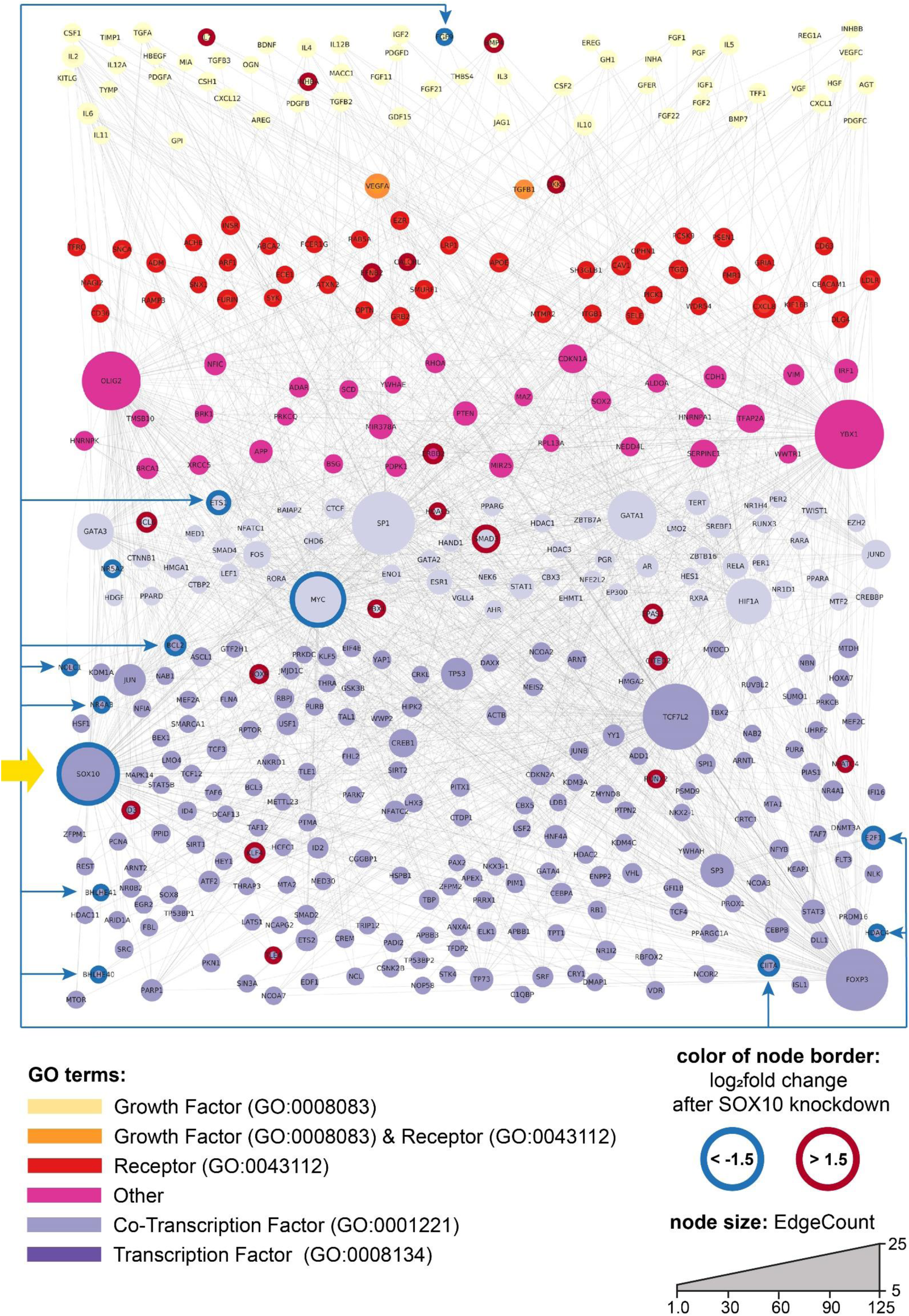
Target selection network. Target selection network for the discovery of new candidates for targeted therapy in UM. The node colors represent different GO terms, and the node size correlates with the EdgeCount of each node. Data obtained from the RNA sequencing analysis after SOX10 knockdown experiments in 92.1 and Mel270 were implemented to color-code the alterations in gene expression of the network members. The colors of the outer ring border of each node show if the gene of interest was downregulated (log_2_fold change ≤ 1.5, blue) or upregulated (log_2_fold change > 1.5, red) upon SOX10 inhibition. Blue arrows: positions of downregulated genes, i.e. potential targets. Yellow arrow: position of SOX10 within the network.

### SOX10 knockdown affects E2F expression and E2F inhibition triggers S phase arrest and apoptosis

Using this target selection network, we searched for entities in the network that were downregulated upon SOX10 inhibition and whose function can be modulated by either commercially available small molecule inhibitors or blocking antibodies, as we thought that inhibiting these candidates may induce similar pro-apoptotic effects as direct siRNA-mediated SOX10 inhibition. Additionally, we were particularly interested in proteins involved in cell cycle regulation or apoptosis. The transcription factor E2F1, which is crucially involved in the transition from G1 to S phase ^49^, met these criteria and was selected as the first suitable candidate for pharmaceutical inhibition in an *in vitro* proof-of-concept study. E2F1 was significantly downregulated in both 92.1 and Mel270 cells after SOX10 knockdown according to the RNA sequencing analysis, which was confirmed by qPCR and immunoblot analyses (Figure 6A, B, C; Figure S8A, B). E2F1 belongs to a family of eight closely related E2F transcription factors, which share highly similar DNA binding domains ^50,51^. Thus, we also analyzed the expression levels of other E2F family members that were not initially part of the target selection network and found that almost all of them were significantly decreased in SOX10 knockdown cells (Figure S8C). E2F transcription factor function can be inhibited by small molecule pan-E2F inhibitor (E2Fi) HLM006474, which has also shown potent antitumor efficacy in CM ^52,53^. HLM006474 incubation for 96 h massively decreased cell viability in all tested UM cells in a dose-dependent manner (Figure 6D; Figure S8D). Propidium iodide staining revealed a significantly increasing number of cells in the S phase of the cell cycle (Figure 6E), Rb hypophosphorylation, and downregulation of cell cycle-regulating proteins like cyclin E1, CDK2, cyclin A2, and cyclin D1 (Figure 6F; Figure S9), indicating S phase arrest induction. As E2F1 promotes its own gene expression in a positive auto-feedback loop through E2F1 binding sites in its promoter region ^54^, HLM006474 treatment also decreased E2F1 protein expression levels as expected. Flow cytometry analyses demonstrated an increase in annexin V-positive cells (Figure 6G), an increased cleavage of caspase 9, caspase 3, and PARP, and a decreased Bcl-2 expression, which is typically for apoptotic cells. Delayed DNA synthesis due to decreased expression of S phase can cause replicative stress including collapsing replication forks that may lead to DNA damage ^55^. Thus, we analyzed γ-

**Figure 6:**
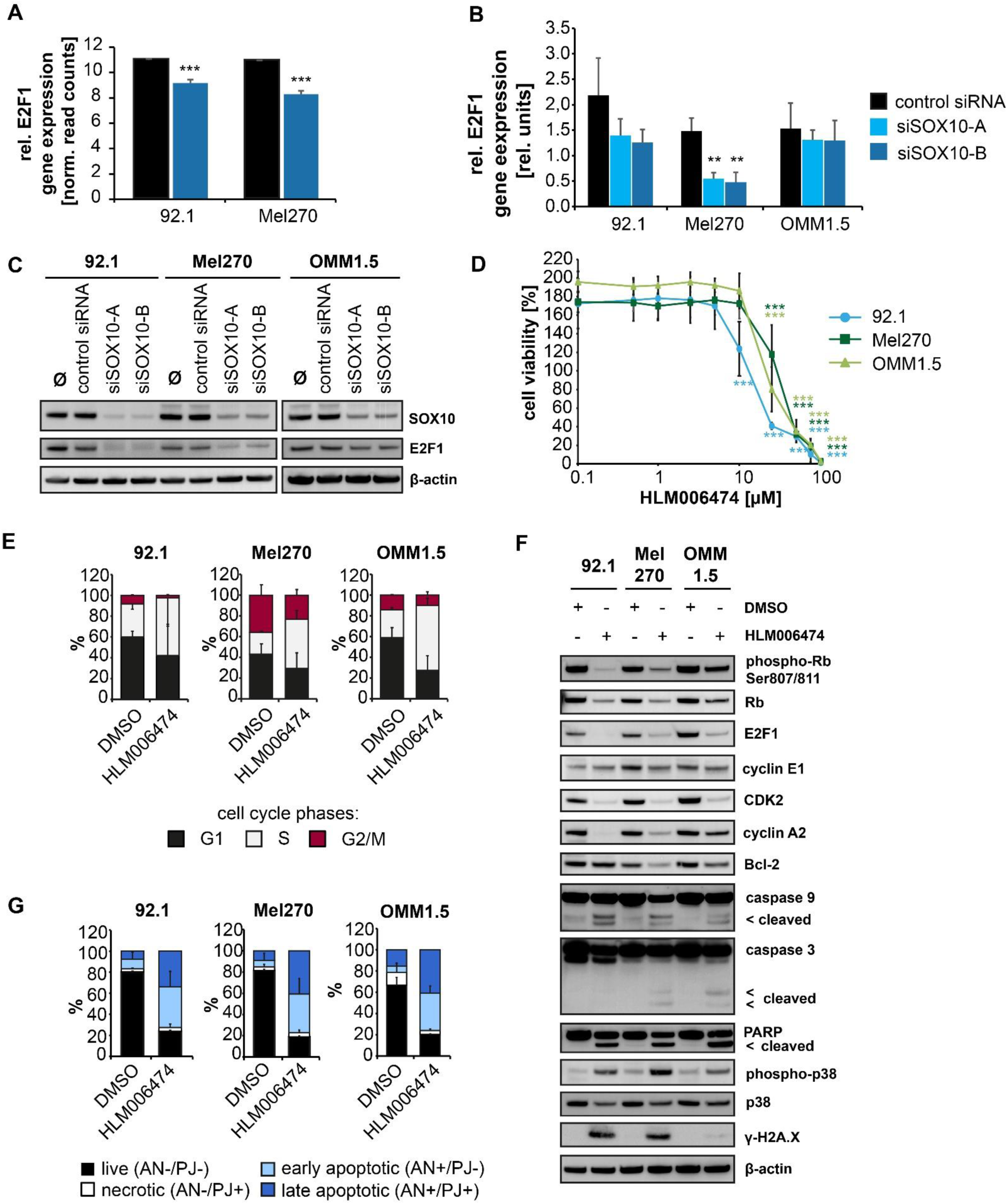
E2F transcription factors are putative new candidates for targeted therapy in UM. (A) Relative E2F1 gene expression in 92.1 and Mel270 transfected with control siRNA or siSOX10-B for 24 h, determined by RNA sequencing, normalized read counts, mean ± SD, n = 4. ***: p < 0.001 vs. control siRNA. (B) Relative E2F1 mRNA expression levels in 92.1, Mel270, and OMM1.5, normalized to GAPDH expression. Relative expression level of 1205Lu was set to 1, data represents mean ± SD, n=3. **: p < 0.01 vs. control siRNA. (C) E2F1 and SOX10 protein expression in 92.1, Mel270, and OMM1.5 48 h after transfection with control siRNA, siSOX10-A, or siSOX10-B. Ø: cells treated with transfection reagent Lipofectamine RNAiMAX only. β-actin served as loading control. (D) Cell viability of 92.1, Mel270, and OMM1.5 incubated with pan-E2Fi HLM006474 or DMSO (control, 100%) for 96 h, determined by CellTiter-Blue® Cell Viability Assay, mean ± SD, n = 3. ***: p < 0.001 vs. DMSO. (E) Cell cycle analysis in UM cell lines 92.1, Mel270, and OMM1.5 incubated with 50 µM HLM006474 or DMSO (control) for 72 h, mean ± SD, n = 3. (F) Protein expression of phosphorylated Rb (phospho-Rb) at Ser807/811, Rb, E2F1, cyclin E1, CDK2, cyclin A2, Bcl-2, caspase 9 and 3, PARP (including cleaved caspase and PARP fragments), phosphorylated p38 (phospho-p38), p38, and histone variant H2A.X phosphorylated at Serin 139 (γ-H2A.X) in 92.1, Mel270, and OMM1.5 incubated with 50 µM HLM006474 or DMSO (control) for 96 h. β-actin served as loading control. (G) Results of annexin V (AN) -propidium iodide (PI) staining and flow cytometry in UM cell lines 92.1, Mel270, OMM1.5 incubated with 50 µM HLM006474 or DMSO (control) for 96 h representing living (AN-/PI-), necrotic (AN-/PI+) and AN+ early and late apoptotic cells. Mean ± SD, n = 3.

H2A.X expression levels and observed an increase in HLM006474-treated cells. Additionally, an increase in p38 phosphorylation was detected, indicating an activation of p38 stress kinase signaling. Similar to SOX10 and MITF inhibition, p53 expression levels did not change upon HLM006474 treatment, suggesting that the effects were mediated in a p53-independent manner. To substantiate the molecular mechanism linking SOX10 and E2F1 to apoptosis in UM, we derived a subnetwork with genes that appear in our UM network and either are connected to E2F1 and SOX10 or are transcriptional targets of any of these transcription factors. We used GO analysis to determine if the genes were associated to positive or negative regulation of apoptosis, and projected the fold change expression of the genes in the SOX10 knockdown experiment. The network suggests multiple mechanisms by which SOX10 inhibition can promote apoptosis with or without E2F1 downregulation, including MITF (Figure 7; Figure S10). Further, several E2F1 apoptosis-related targets were differentially regulated. Finally, we found several potential molecular pathways through which SOX10 inhibition may promote E2F1 downregulation, as e.g. the axis SOX10-OLIG2-NR4A1-E2F1 or SOX10-TCF7L2-NR4A1-E2F1 which may be explored in detail in future studies. In summary, using this computational network-based approach, we identified the E2F transcription factors as putative candidates for targeted therapy in UM. SOX10 inhibition diminished E2F expression levels, and pan-E2Fi HLM006474 demonstrated strong cytotoxic effects in all tested UM cell lines accompanied by an S phase arrest, p38 stress kinase activation, and apoptosis already at low micromolar concentrations. These data indicate that HLM006474 may be a promising drug candidate for further *in vivo* studies in UM and highlight the potential of this computational drug selection approach for the development of new treatment strategies in UM.

**Figure 7:**
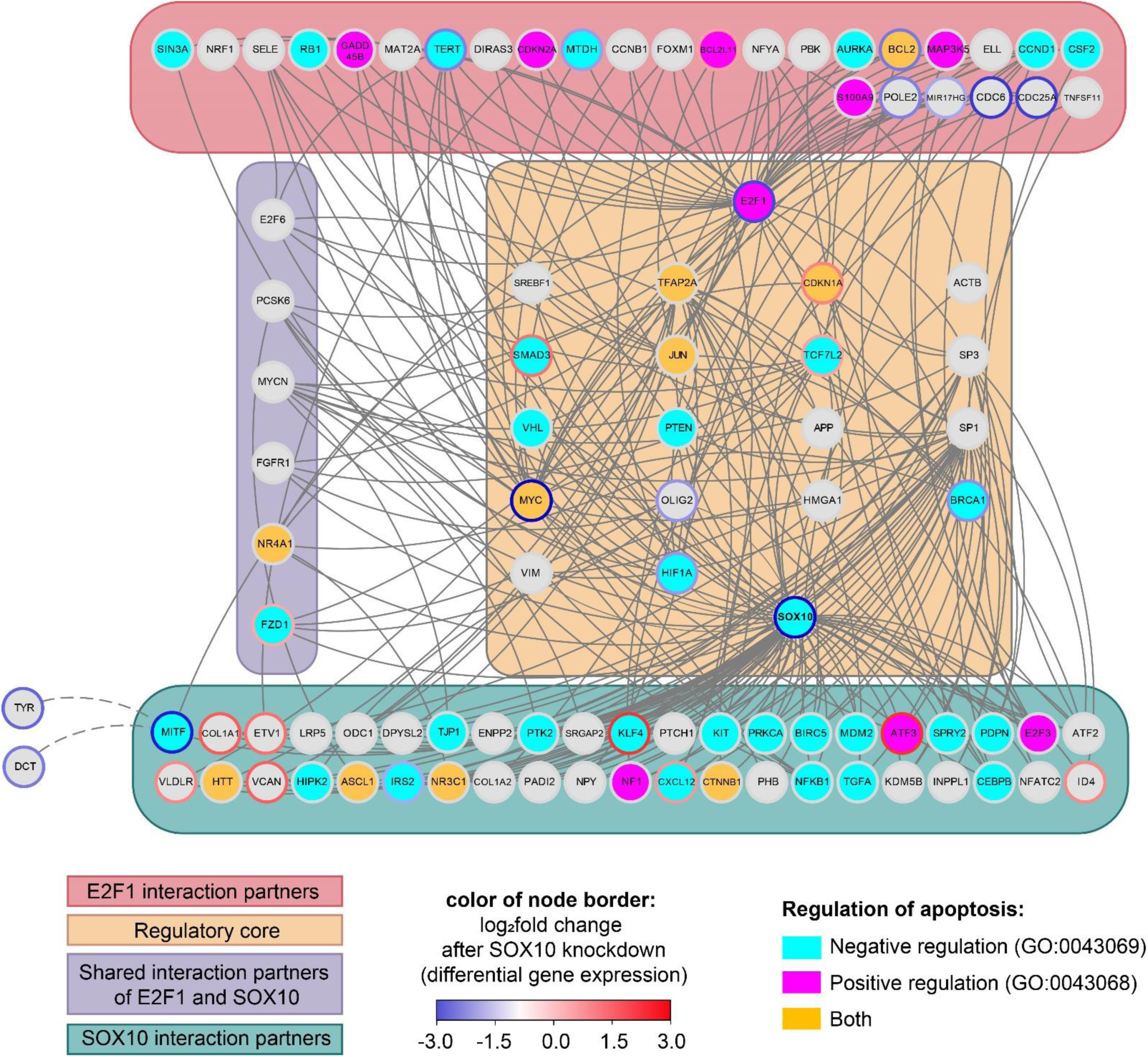
Links of SOX10 and E2F1 to apoptosis in UM. UM subnetwork with genes that are connected to E2F1, SOX10, or transcriptional targets of any of these transcription factors. The nodes were colored according to their occurrence in the Gene Ontology annotations for negative regulation of programmed cell death (GO:0043069, cyan), positive regulation of programmed cell death (GO:0043068, pink), or both (orange). The node borders were colored utilizing the log_2_fold change in gene expression obtained from the RNA sequencing analysis after SOX10 knockdown. Nodes were grouped by either their occurrence in the regulatory core network (light orange), their properties as interaction partners of SOX10 (light green), E2F1 (light red), or both (light purple). Two nodes for MITF-regulated direct target genes TYR and DCT were added manually and highlighted with a dashed connection to MITF.

## Discussion

Despite the massive progress that has been made in recent years regarding the treatment of CM, including the introduction of BRAF and MEK inhibitors and ICB, the clinical management of UM and particularly its advanced stages is still challenging, as most of these new therapy options are hardly effective in UM ^5^. Additionally, the tumor mutational burden in UM is relatively low in contrast to e.g. CM, and the few recurrently mutated genes in UM like *GNAQ* or *GNA11* cannot be targeted therapeutically yet ^1,6^. Thus, new ways must be explored to find effective and specific new therapy options for the patients. In the past, several NC-related transcription factors have been identified to contribute to cancer initiation and progression in CM ^20^; however, it remained unclear until now if they also play a role in UM and if this knowledge can be exploited for the development of new therapy options in UM. In this work, we found high SOX10 expression in UM in contrast to other NC-related transcription factors such as POU4F1 and MSX1 and demonstrated that the SOX10-MITF axis is particularly important for UM survival, as its inhibition impaired UM cell viability, induced a G1 arrest, and massively triggered apoptosis. As neither SOX10 nor MITF can be targeted by small molecule inhibitors to date, we used an innovative UM-specific computational target selection strategy to identify more druggable SOX10-associated proteins whose function can be blocked instead of SOX10 or MITF. To test this approach, we identified and confirmed the E2F transcription factor family as potential druggable targets in UM in a proof-of-concept *in vitro* study by using pan-E2F inhibitor HLM006474, resulting in cell cycle arrest and apoptosis in all tested UM cell lines.

SOX10 protein expression has been explored in UM primary tumors before ^27,28^, but not yet in metastases. According to our data, SOX10 is widely expressed in both primary tumors and metastases; however, UM primary tumor-derived cell lines 92.1 and Mel270 were more susceptible to SOX10 inhibition than UM cell line OMM1.5 which was initially derived from a liver metastasis of the same patient as Mel270 ^41^, or CM-derived cell line 1205Lu which was selected in nude mice for lung metastasis formation after subcutaneous injection of CM cell line WM793 ^56^. Nevertheless, OMM1.5 and 1205Lu were not completely resistant to SOX10 knockdown as cell viability still decreased, albeit with a delay. Further studies have to elucidate if SOX10 truly plays a more important role in primary tumors than in metastases. The results of this study in UM are mostly in line with previous findings in CM, which show that SOX10 is essential for cell survival and cell cycle progression ^25,26^, indicating that the role of SOX10 is similar in UM and CM despite the fundamental differences regarding the tumor biology of these melanoma subtypes.

MITF is the master regulator of melanogenesis and a well-characterized target gene of SOX10 ^47^. We observed that MITF expression correlated with SOX10 expression in UM, and several of the SOX10 knockdown-related effects were mediated by MITF. Ectopic MITF expression mitigated the pro-apoptotic effects of SOX10 inhibition; however, it was not able to completely abolish the cytotoxic effects of SOX10 loss. Similar to SOX10 knockdown, the decrease in cell viability of OMM1.5 after siMITF transfection was also delayed compared to 92.1 and Mel270 ^57^. Together, these results indicate that MITF also has a pro-survival function in UM. Interestingly, these results are contradictory to other findings in UM, which show that MITF expression is associated with proliferative activity in UM ^58^, and loss of mitfa expression accelerated GNAQ^Q209L^-driven tumorigenesis in a zebrafish model ^59^. However, given the complex and dynamic functions of MITF in CM heterogeneity and cellular plasticity ^47,60^, and the contradictory findings in UM, the functional role of MITF in UM might be as dynamic and heterogeneous as in CM, indicating that its function may vary depending on its expression levels and expressed cofactors.

One of the main goals of this work was to translate these findings into novel therapeutic approaches by taking advantage of the crucial role of SOX10 for UM survival. Therefore, we applied a computational UM-specific network-based target selection strategy to identify druggable proteins that can be inhibited by small molecule inhibitors instead of SOX10 or MITF and may serve as candidates for new targeted therapies in UM. Following this approach, we selected the E2F transcription factors as their expression levels were influenced by SOX10, and demonstrated that treatment with pan-E2Fi HLM006474 significantly reduced cell viability and resulted in cell cycle arrest in the S phase, increased γ-H2A.X expression, p38 activation, and apoptosis in all tested UM cell lines. Notably, E2F1 and HLM006474 were initially selected based on the target selection network as putative target and corresponding small molecule inhibitor, respectively; however, HLM006474 is not specific for E2F1 but inhibits other members of the E2F transcription factor family as well ^52^. Unfortunately, no selective E2F1 inhibitor is currently available as far as we know. Nevertheless, the lack of HLM006474 specificity may not necessarily be a disadvantage as previous experiments have shown that E2F transcription factors can step in to compensate for the loss of other E2F family members due to their homology, at least in short-term studies ^50,51^. Thus, using a pan-E2Fi may exert a better treatment efficacy as an inhibitor selectively targeting E2F1, as bypassing E2F1 inhibition is not possible. Furthermore, the loss of SOX10 decreased the expression of almost all E2F transcription factor family members, suggesting that downregulation upon SOX10 knockdown is not restricted to E2F1 alone. Therefore, these results may be also interpreted as HLM006474 mimicking SOX10’s pan-E2Fi function as it prevents the compensatory role of other E2F family members as well.

Achieving good antitumor efficacy with limited toxic effects on non-tumor tissue is one of the main goals of systemic cancer therapy. Noteworthy, SOX10 knockdown did not affect the cellular morphology or cell viability of HM, indicating that normal cells can compensate for SOX10 loss in contrast to tumor cells. However, to our knowledge, no specific small molecule inhibitor of SOX10 exists; thus, other ways of exploiting the effects of SOX10 inhibition for therapeutic purposes are needed. In this study, we identified the E2F transcription factors as potential candidates for targeted therapy that can be inhibited instead of SOX10 and pan-E2Fi HLM006474 as the corresponding drug candidate. However, HLM006474 has not been tested in humans yet, and no clinical trials testing this compound are currently registered at ClinicalTrials.gov. Additionally, E2F1 in particular is widely expressed in the human body ^61^, bearing the risk for systemic side effects of HLM006474 treatment. However, no cytotoxic effects of HLM006474 on normal cells such as fibroblasts and keratinocytes have been observed so far in preclinical *in vitro* studies ^52,53^. Furthermore, CM-bearing mice treated with HLM006474 did not show any signs of toxicity ^53^. These findings suggest that HLM006474 may be a promising drug candidate for UM treatment, although E2F1 expression is not restricted to UM cells only. Nevertheless, further preclinical studies are needed to explore the efficacy and safety profiles of HLM006474.

In summary, SOX10 is widely expressed in UM, and the SOX10-MITF axis is crucial for UM cell cycle progression and cell survival. Using a network-based strategy, the E2F transcription factors were identified as druggable SOX10-associated proteins that can be inhibited by small molecule pan-E2Fi HLM006474 instead of SOX10 or MITF, and HLM006474 also triggered apoptosis in UM cell lines similarly to siRNA-mediated SOX10 knockdown. The successful combination of computational approaches and *in vitro* experiments in this study demonstrates a new way of translating basic knowledge about transcription factor biology into developing new treatment approaches and may be an attractive approach, especially for rare and difficult-to-treat tumor entities.

## Acknowledgements

We thank the dermatohistopathology core, Waltraud Fröhlich, Annett Hamann, and Lena Stich (Department of Dermatology, Uniklinikum Erlangen) for excellent technical assistance. Furthermore, we thank Klaus Griewank (University Hospital Essen, Germany), Martine Jager (University of Leiden, The Netherlands) and Meenhard Herlyn (The Wistar Institute, Philadelphia, USA) for kindly providing the cell lines.

## Data availability statement

RNA seq data from the SOX10 knockdown experiments in UM cell lines 92.1 and Mel270 generated in this study are publicly available in the Gene expression omnibus (GEO) database (GEO ID: GSE261687, accessible via https://www.ncbi.nlm.nih.gov/geo/query/acc.cgi?acc=GSE261687). Anonymized patient data are available from the corresponding author upon reasonable request. All other data are available in the main text or the Additional files.

## Ethics approval statement and patient consent statement

The studies involving human subjects were reviewed and approved by the Ethics Committee of the Friedrich-Alexander Universität Erlangen-Nürnberg. The principal investigator and all other investigators ensured that the studies were conducted in full conformity with the principles set forth in the Declaration of Helsinki. Informed written consent was obtained from each patient before sample collection.

## Author contributions

Conceptualization: An.W., C.B., M.V.H., Formal Analysis: An.W., C.L., Ad.W., E.G., E.A.T.K., Funding Acquisition: An.W., C.B., M.V.H., B.S.T., Investigation: An.W., C.K., C.L., Ad.W., E.G., Methodology: A.W., C.K., C.L., Ad.W., J.V., M.V.H., Project Administration: J.V., C.B., M.V.H., Resources: J.D., B.S.T., Mi.E., C.V., S.S., J.N., Ma.E., Software: C.L., Ad.W., X.L., J.V., Supervision: C.B., M.V.H., Visualization: An.W., C.L. Ad. W., E.G., X.L., Writing – Original Draft Preparation: An.W., M.V.H., Writing – Review & Editing: C.K., C.L., Ad.W., X.L, E.G., E.A.T.K., J.D., B.S.T., Mi.E., C.V., S.S., J.N., Ma.E., J.V., C.B. All authors read and approved the final version of the manuscript and were responsible for the decision to submit the manuscript.

## Additional files

Additional file A1: Supplementary Materials and Methods

Additional file A2: Supplementary Figures

Additional file A3: Supplementary Tables

## SUPPLEMENTARY MATERIALS *AND* METHODS

### RNA sequencing of UM tumor samples

Gene expression levels of *SOX10, MSX1, POU4F1*, and *SOX9* were determined in UM tumor specimens obtained from two independent patient cohorts via RNA sequencing. The tumor samples of the first cohort included 12 primary tumors and three liver metastases of UM patients from the Uniklinikum Erlangen, which were analyzed using the Nanopore® sequencing technology. Sequencing libraries were prepared with the Direct RNA Sequencing kit or the PCR-cDNA Sequencing kit (both from Oxford Nanopore Technologies, Oxford, UK) depending on the amount of available RNA following the manufacturer’s instructions. Sequencing was carried out on R9.4 flow cells using a MK1B-MinION sequencer (Oxford Nanopore Technologies), and RNA sequencing data was analyzed as described previously ^1^. The tumor samples of the second independent cohort included 16 metastases (liver: n=14, breast: n=1, lung: n=1) from UM patients who participated in a phase 1 clinical trial investigating IKKb-matured, RNA-loaded dendritic cells for the treatment of metastasized UM at the Uniklinikum Erlangen (NCT04335890) ^2^. Library preparation, RNA sequencing, and data analysis were performed by a commercial provider (CeGaT, Tübingen, Germany) using the KAPA RNA HyperPrep Kit with RiboErase (Roche Diagnostics, Mannheim, Germany) for library preparations and a NovaSeq™ 6000 instrument for sequencing (Illumina, San Diego, California, USA).

### Immunohistochemical staining of SOX10 in UM metastases

To investigate SOX10 protein expression in UM metastases, 5 µm sections of formalin-fixed and paraffin-embedded tumor samples were stained using anti-SOX-10 antibody (EP268, Cell Marque, Rocklin, California, USA; RRID:AB_2941085) at a 1:100 dilution in Dako REAL Antibody Diluent (Agilent Technologies, Santa Clara, California, USA). Detection was performed by the alkaline phosphatase method using the ultraView Universal Alkaline Phosphatase Red Detection Kit and Hematoxylin II (both from Roche Diagnostics) for counterstaining of nuclei according to the manufacturer’s instructions. Microscopic images were taken with a BZ-X800 microscope (Keyence, Neu-Isenburg, Germany) and analyzed with the image analysis software QuPath ^3^.

### Cell culture

UM cell lines were cultivated in RPMI1640 medium supplemented with 2 mM L-glutamine (Gibco by Life Technologies, Waltham, Massachusetts, USA), 10% fetal bovine serum (Merck, Darmstadt, Germany), and 1x antibiotic-antimycotic (Invitrogen, Waltham, Massachusetts, USA). CM cell line 1205Lu was cultivated in tumor 2% medium as described previously ^4^. Human epidermal neonatal melanocytes (HM) were purchased from Cell Systems Biotechnologie Vertrieb (Troisdorf, Germany) and cultivated in DermaLife® Basal Medium supplemented with DermaLife® M LifeFactors kit (LifeLine Cell Technology, Frederick, Maryland, USA) and 1x antibiotic-antimycotic for a maximum of 10 passages. All cells were maintained in a humidified incubator at 37°C and 5% CO_2_ atmosphere and regularly tested for mycoplasma contamination using the Venor®GeM Classic Mycoplasma Detection Kit for conventional PCR (Minerva Biolabs, Berlin, Germany) according to the manufacturer’s instructions.

### RNA isolation, cDNA synthesis, and quantitative real-time-PCR (qPCR) analysis

Total RNA was extracted using the RNeasy® Mini Kit (Qiagen, Hilden, Germany) following the manufacturer’s instructions. RNA concentration was determined on a Nanodrop 2000c spectrophotometer (Thermo Fisher Scientific). One µg of RNA was transcribed using the Expand™ Reverse Transcriptase Kit (Roche, Penzberg, Germany) and oligo(dT)-oligonucleotide primers (Eurofins Genomics, Ebersberg, Germany). Oligonucleotide primers for SOX10, MITF, and GAPDH were designed with the software Assay Design Center (Roche; https://lifescience.roche.com/en_de/brands/universal-probe-library.html#assay-design-center) and E2F1 primers were designed with the software Primer-BLAST ^5^. The sequences are presented in Supplementary Table 4. The LightCycler® TaqMan® Master Kit (Roche), 200 nM of forward and reverse primers, 100 nM primer-matched hydrolysis probes from the Universal ProbeLibrary set human (Roche), and 50 ng cDNA was used for qPCR which was carried out on a Lightcycler 2.0 instrument (Roche). E2F1 mRNA expression levels were determined using 2x innuMIX qPCR DSGreen Standard master mix (AJ Innuscreen GmbH, Berlin, Germany), 200 nM of forward and reverse primers, and 50 ng cDNA on a qTower^3^ instrument (Analytik Jena, Jena, Germany). Gene expression of untreated 1205Lu cells was set to 1 and gene expression was normalized to glyceraldehyde 3-phosphate dehydrogenase expression (GAPDH).

### Detection of protein expression by immunoblot

Protein was extracted from cells using cell lysis buffer containing 50 mM Tris, 250 mM NaCl, 1 mM EDTA, 0.1 % triton X-100 (all from Sigma-Aldrich), cOmplete™ Mini protease inhibitor cocktail, and PhosSTOP™ phosphatase inhibitor cocktail (both from Roche). Protein concentration was determined with the Pierce™ BCA Protein Assay kit on a Nanodrop 2000c spectrophotometer (both from Thermo Fisher Scientific). Gel electrophoresis using 4-12% gradient polyacrylamide gels in MES SDS running buffer and blotting on PVDF membranes was performed with XCell SureLock™ Mini-Cells and the NuPAGE Novex System (all from Invitrogen) following the manufacturer’s instructions. Membranes were blocked with 1x Western Blocking Reagent (Roche) in phosphate-buffered saline (PBS) and 45.5 mM NaF to prevent unspecific antibody binding and incubated with primary antibodies (Supplementary Table 5) overnight at 4°C while gently shaking. After three washing cycles with 0.1% Tween-20/PBS, the membranes were incubated with horseradish peroxidase (HRP)-linked secondary antibodies (Supplementary Table 6) for 1 h at room temperature. After another three washing cycles, membranes were incubated with Amersham™ ECL Prime Western Blotting Detection Reagent (GE Healthcare, Chalfont, United Kingdom) for 5 min, and chemiluminescence detection was performed with Amersham™ Hyperfilm™ ECL X-ray films (GE Healthcare) and an RP X-OMAT developer (Kodak, Stuttgart, Germany) or an ImageQuant LAS4000 camera-based detection system (GE Healthcare). For sequential probing of immunoblots after chemiluminescence detection, membranes were washed in 0.1% Tween-20/PBS and incubated in blocking solution supplemented with 0.02% NaN_3_ (all from Sigma-Aldrich) for 1 h at room temperature to destroy HRP function as published by Kaufmann ^6^, and rehybridized with primary antibodies as described above.

### Caspase 3 inhibition experiments

UM cell lines 92.1 and Mel270 were seeded in 6-well plates and transfected with SOX10-specific siRNAs or control siRNA on the following day for 48 h as described in the materials and methods section. Cells were preincubated for 1 h before transfection and during the whole transfection experiment with the irreversible caspase 3 inhibitor Z-DEVD-FMK (Selleck Chemicals LLC, Houston, Texas, USA) or equal amounts of DMSO (Sigma-Aldrich, Taufkirchen, Germany) as control. Cell viability was determined by the CellTiter-Blue® Cell Viability Assay (Promega, Madison, Wisconsin, USA) according to the manufacturer’s protocol. The fluorescence intensity of the supernatant (excitation: 530 nm, emission: 590 nm) was measured in a plate reader Glo-Max® Explorer plate reader (Promega) and relative cell viability compared to respective control cells was calculated in Microsoft Excel. Cleavage of caspase 3 and PARP was determined using immunoblots as described above.

### SOX10 knockdown rescue experiments

For rescue experiments, Mel270 cells seeded in 6-well plates the day before were transfected with 2 µl P3000 reagent, 3.75 µl Lipofectamine™ 3000 Reagent (both from Invitrogen, Waltham, Massachusetts, USA), and 2.5 µg plasmid DNA per well (pCMV6-MITF-Myc-DDK expression plasmid with open reading frame for Myc- and FLAG® (DDK)-tagged melanocytic isoform of MITF (MITF transcript variant 4) (# RC209561, Origene, Rockville, Maryland, USA) or pCMV6-Entry vector as control (# PS100001, Origene)). The day following plasmid transfection, cells were transfected again with siSOX10-B or control siRNA as in the materials and methods section and incubated in a humidified incubator at 37°C and 5% CO_2_ atmosphere for 24 h and 48 h.

### Analysis of transcriptomic changes in SOX10 knockdown cells

RNA sequencing analysis of pooled total RNA derived from four individual SOX10 knockdown experiments using siSOX10-B or control siRNA was performed by a commercial provider (CeGaT, Tübingen, Germany) using the KAPA RNA HyperPrep Kit with RiboErase (Roche) for library preparation and a NovaSeq™ 6000 instrument (Illumina, San Diego, California, USA) for sequencing. RNA quality was checked on a Bioanalyzer 2100 (Agilent Technologies, Santa Clara, California, USA) before RNA sequencing. Further analyses were carried out in R Studio using the packages DESeq2, AnnotationDbi, pheatmap, and ggplot2. Obtained FASTQ reads were mapped to the human reference genome assembly GRCh38.p2 (hg38)) in R with Gencode Annotation (version 22) ^7^ and short read aligner STAR (version 2.0201) ^8^. Differences in the gene expression of SOX10 knockdown and control cells with a family-wise error rate of > 0.05 were considered significantly differentially expressed. Putative SOX10 target genes were identified by comparing the data with SOX10 binding data (10,000 bp upstream to 5,000 bp downstream of transcription start sites) obtained from SOX10 chromatin immunoprecipitation and sequencing (ChIP seq) experiments in melanocytes ^9^. Gene set enrichment analysis was performed using Enrichr with the MSigDB hallmark gene sets ^10,11^. For SOX10, we used the differentially expressed genes (n = 1,949) as the input gene list and, for MITF, we used its targets that were differentially expressed in the SOX10 knockdown condition (n = 435) based on a list of MITF target genes (n = 2552) collected from TRANSFAC and data from the literature^12, 14^ The annotated genes in the entire genome were used as the background gene list. Gene sets with an adjusted p-value ≤ 0.05 were considered to be significant and computed adjusted p-values computed using the false discovery rate approach. The results were visualized as heat maps and volcano plots using ComplexHeatmap and ggplot2, respectively ^15^.

### Reconstruction of a SOX10-based UM-specific protein interaction network: Calculation of weighted ranking score

To detect the most important nodes and their interactions, a weighted ranking score of the previously identified motifs was calculated, which is comparable to the one used by Khan et al. ^16^:

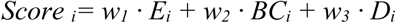

Where each motif i is calculated with the different weight settings w_1_, w_2_, and w_3_ that define the importance of the three ranking factors: E_i_ is the respective expression value across the nodes forming the motif i. BC_i_ is the average betweenness centrality across the nodes forming the motif i. D_i_ is the average node degree across the nodes forming the motif i. Similar to Khan et al. ^16^ and Cantone et al. ^17^, the weighting factors sum up to 1, where w_1_ was fixed to 0.5 to underline the importance of motif expression. The values of w_2_ were set from 0.05 up to 0.45 in 0.05 iterative steps and the values of w_3_ result from the calculation w_3_ = 0.5 – w_2_. The motif scores of each motif i were calculated for each combination of weighting factors and the different scores of the same motifs were pareto-optimized with the psel method using the R package rPref (version 1.3) ^18^. The expression values were derived from the 80 pooled primary UM samples previously added in the pruning step described in the materials and methods section and the respective betweenness centrality and degree of a node were used from the Network Analyzer output.

### Development of the target selection network

The underlying core was then used as starting point for creating the target selection network. The core genes and their respective first neighbors were extracted from the comprehensive pruned UM network (Supplementary table 3) and the nodes were filtered and re-annotated according to their basic functionality as classified by the gene ontology (GO) terms for growth factors (GO:0008083), receptors (GO:0043112), co-transcription factors (GO:0001221), and transcription factors (GO:0008143). Self-loops were removed and the resulting network was reformatted using the CerebralWeb implementation and the differential expression values collected from the SOX10 knockdown experiments were projected into the network’s visualization.

### Reconstruction of a SOX10-E2F1 apoptosis connection network in UM

To link SOX10 to the regulation of E2F1 and apoptosis in UM, we used the Cytoscape plugin PathLinker (version 1.4.3) ^19^ to obtain the top 150 undirected shortest paths connecting SOX10 and E2F1 from the full UM network. These shortest paths were merged in a separate sub-network and the nodes were grouped according to their occurrence in the regulatory core network, their properties as interaction partners of SOX10, E2F1, or both. Additionally, the nodes were colored according to their occurrence in the GO annotations for negative regulation of programmed cell death (GO:0043069), positive regulation of programmed cell death (GO:0043068), or both.

## SUPPLEMENTARY FIGURES

**Supplementary Figure S1:**
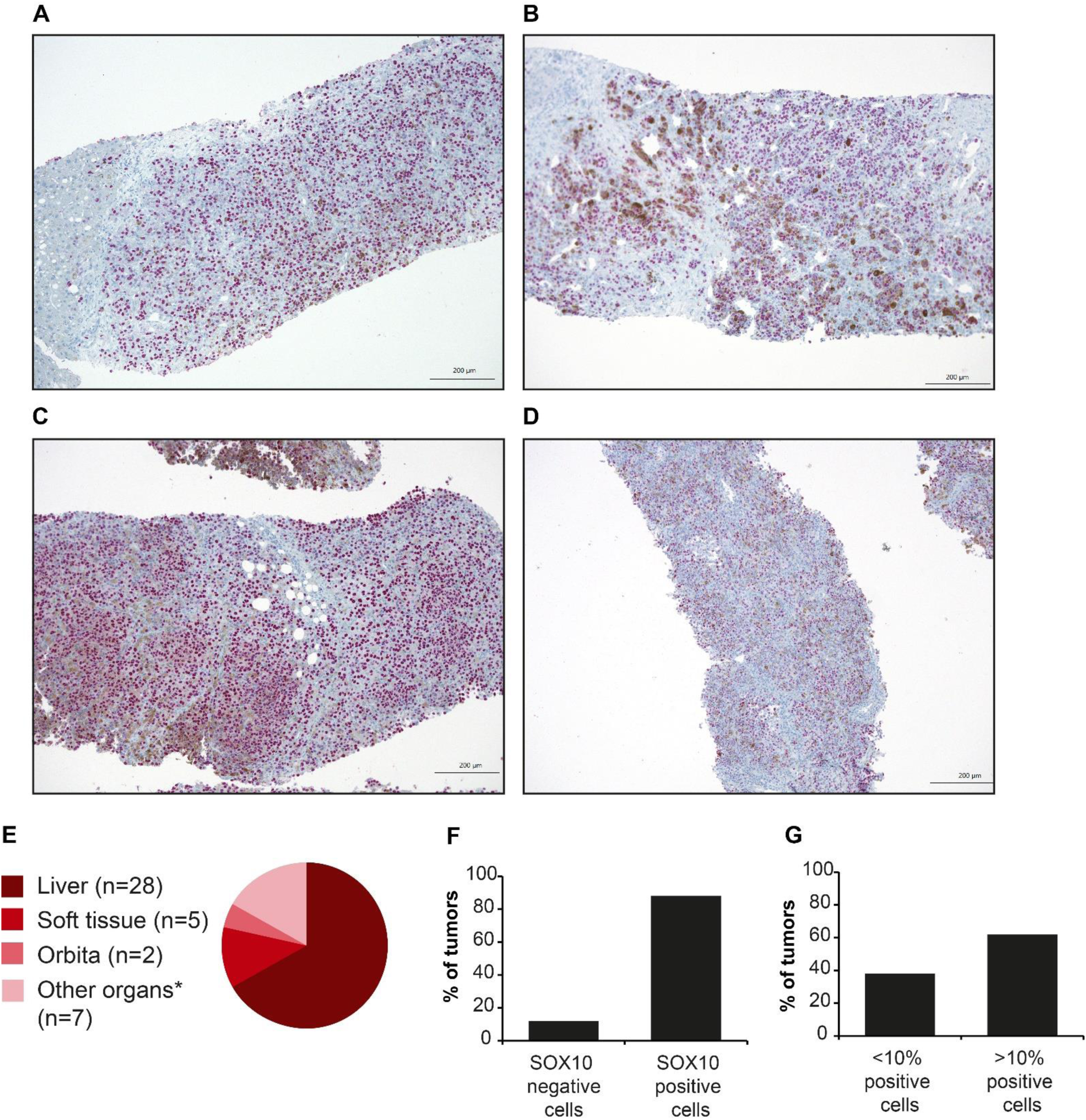
Immunohistochemical staining of SOX10 in UM metastases. Representative microscopic images of immunohistochemical SOX10 staining of UM distant metastases: **(A, B)** liver metastases, **(C)** metastasis of the greater omentum, **(D)** kidney metastasis. **(E)** Localization of analyzed UM metastases investigated for SOX10 protein expression via immunohistochemical staining, n=42. *: greater omentum, ear, breast, lymph node, adrenal glands, kidney, lower back (n=1 each), scale bar = 200 µm. **(F)** Percentage of tumors with SOX10 negative (< 1% SOX10-expressing cells) and SOX10 positive cells (>1% SOX10-expressing cells), n=42. **(G)** Percentage of tumors with < 10% and > 10% SOX10 positive cell, n=42.

**Supplementary Figure S2:**
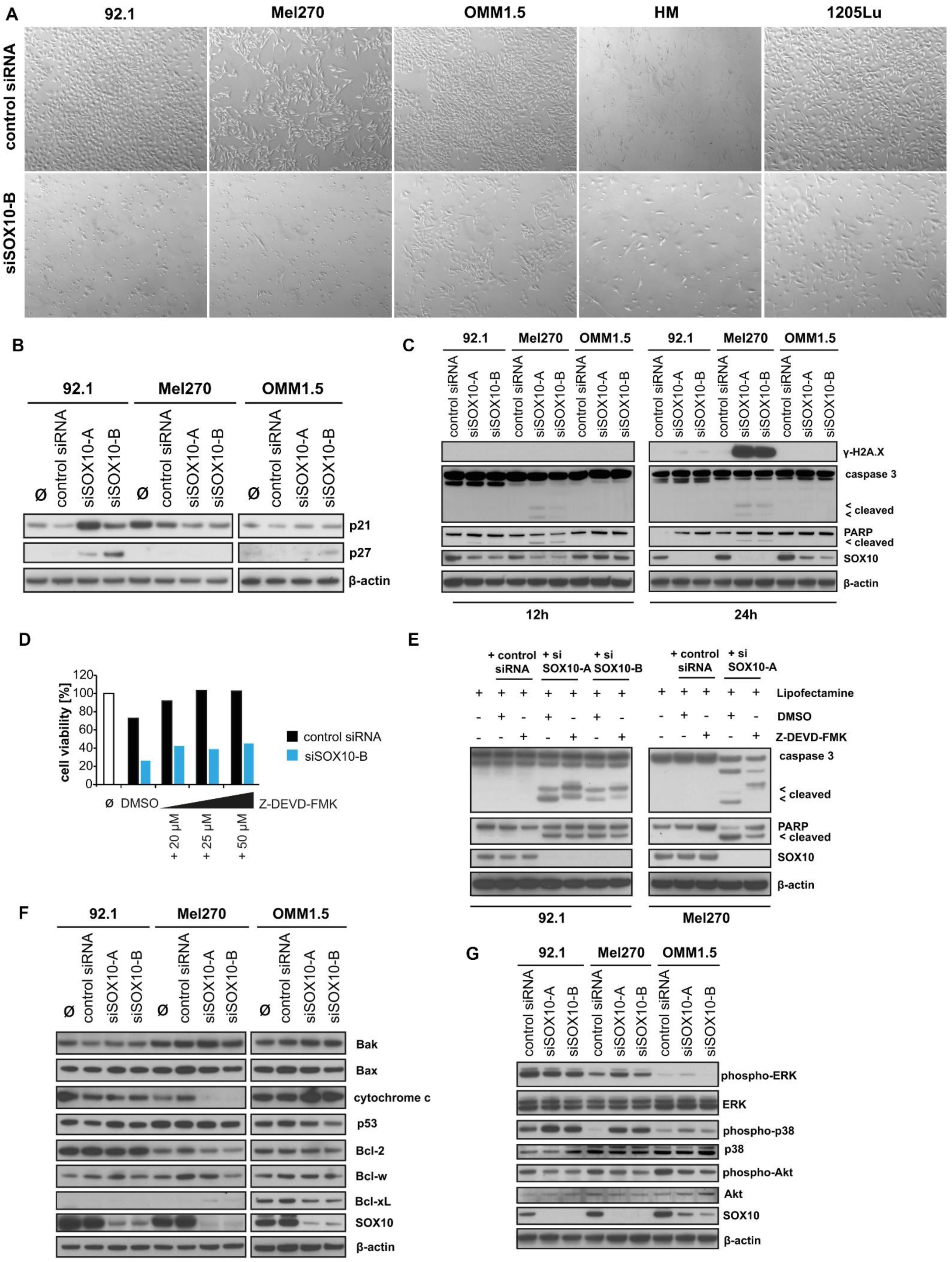
Effects of SOX10 knockdown. **(A)** Microscopic images of UM cell lines 92.1, Mel270, OMM1.5, human melanocytes (HM), and CM cell line 1205Lu transfected with control siRNA or siSOX10-B for 96 h, 50fold magnification. **(B)** Protein expression of p21 and p27 in 92.1, Mel270, and OMM1.5 48 h after transfection with control siRNA, siSOX10-A, or siSOX10-B. Ø: cells treated with transfection reagent Lipofectamine RNAiMAX only. β-actin served as loading control. **(C)** Protein expression of histone variant H2A.X phosphorylated at Serin 139 (γ-H2A.X), caspase 3, PARP (including cleaved protein fragments), and SOX10 in 92.1, Mel270, and OMM1.5 12h and 24 h after transfection with control siRNA, siSOX10-A or siSOX10-B. β-actin served as loading control. **(D)** Cell viability of UM cell line 92.1 transfected with control siRNA or siSOX10-B for 48 h, determined by CellTiter-Blue® Cell Viability Assay, n = 1. Cells were preincubated for 1 h before transfection and during the whole transfection experiment with the irreversible caspase 3 inhibitor Z-DEVD-FMK or DMSO as control. Ø: untreated cells. **(E)** Protein expression of caspase 3, PARP (including cleaved protein fragments), and SOX10 in 92.1, Mel270, and OMM1.5 48 h after transfection with control siRNA, siSOX10-A, or siSOX10-B and concurrent incubation with 20 µM Z-DEVD-FMK or DMSO as control. The cells were preincubated with 20 µM Z-DEVD-FMK or DMSO for 1h prior to transfection. β-actin served as loading control. **(F)** Protein expression of Bak, Bax, cytochrome c, p53, Bcl-2, Bcl-w, Bcl-xL, and SOX10 in 92.1, Mel270, and OMM1.5 48 h after transfection with control siRNA, siSOX10-A, or siSOX10-B. Ø: cells treated with transfection reagent Lipofectamine RNAiMAX only. β-actin served as loading control. **(G)** Protein expression of phospho-ERK, ERK, phospho-p38, p38, phospho-Akt, and Akt in HM, 92.1, Mel270, OMM1.5, and 1205Lu transfected with control siRNA, siSOX10-A, or siSOX10-B for 24 h. Ø: treated with transfection reagents only. β-actin served as loading control for immunoblots.

**Supplementary Figure S3:**
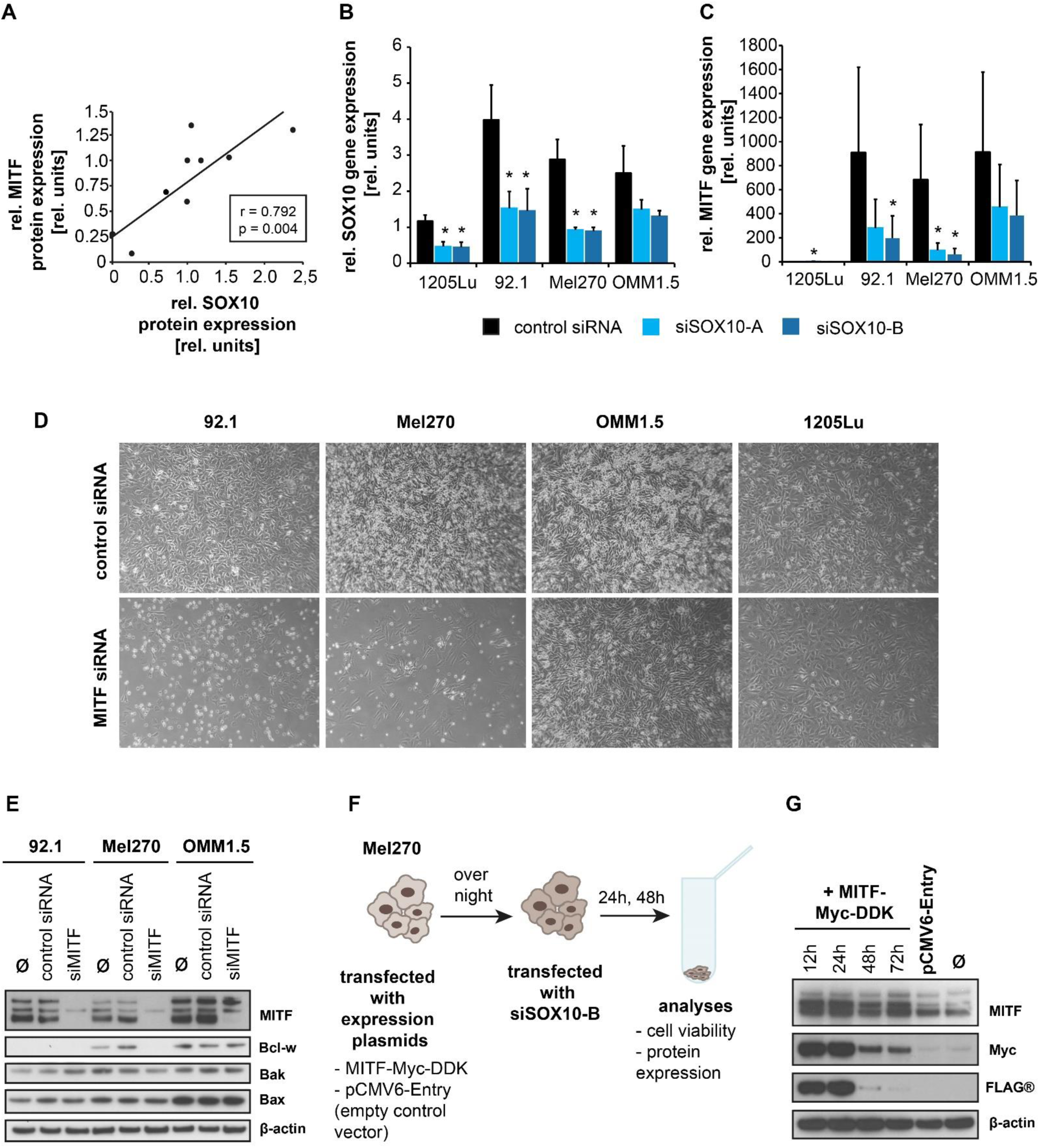
SOX10 target gene MITF contributes to pro-survival effects of SOX10. **(A)** Positive linear correlation between SOX10 and MITF protein expression in UM cell lines and HM, relative expression normalized to β-actin expression. Protein expression levels were determined by densitometric analyses of immunoblots (shown in Fig. 3B) using ImageJ software. Pearson correlation coefficient r = 0.792, p = 0,004. **(B)** Relative SOX10 and **(C)** MITF mRNA expression levels in CM cell line 1205Lu and UM cell lines 92.1, Mel270, and OMM1.5 transfected with control siRNA, siSOX10-A, or siSOX10-B for 24 h, normalized to GAPDH expression. Relative expression level of 1205Lu was set to 1, data represents mean ± SD, n=3. **(D)** Microscopic images of UM cell lines 92.1, Mel270, OMM1.5, and CM cell line 1205Lu transfected with control siRNA or siMITF for 96 h, 50fold magnification. **(E)** Protein expression of MITF, Bcl-w, Bak, and Bax in 92.1, Mel270, and OMM1.5 transfected with control siRNA or siMITF for 24 h. Ø: cells treated with transfection reagent Lipofectamine RNAiMAX only. β-actin served as loading control. **(F)** Scheme depicting the experimental procedure of rescue experiments. Mel270 cells were transfected with either a MITF expression vector (MITF-Myc-DDK), leading to the ectopic expression of a FLAG® (DDK)- and Myc-tagged MITF protein or an empty control vector (pCMV6-Entry), followed by siSOX10-B transfection on the next day. Cell viability analyses and harvest for protein expression analyses were performed 24 h and 48 h after siRNA transfection. **(G)** Time course analysis of ectopic MITF protein expression and Myc and FLAG® tagged protein. Mel270 cells were transfected for 12 h, 24 h, 48 h, and 72h with an MITF expression vector (MITF-Myc-DDK), leading to the ectopic expression of a FLAG® (DDK)- and Myc-tagged MITF protein, confirming ectopic MITF expression during the entire time period of the rescue experiments (=72 h). Controls: transfection with empty vector pCMV6-Entry for 72 h, Ø: untreated cells. β-actin served as loading control.

**Supplementary Figure S4:**
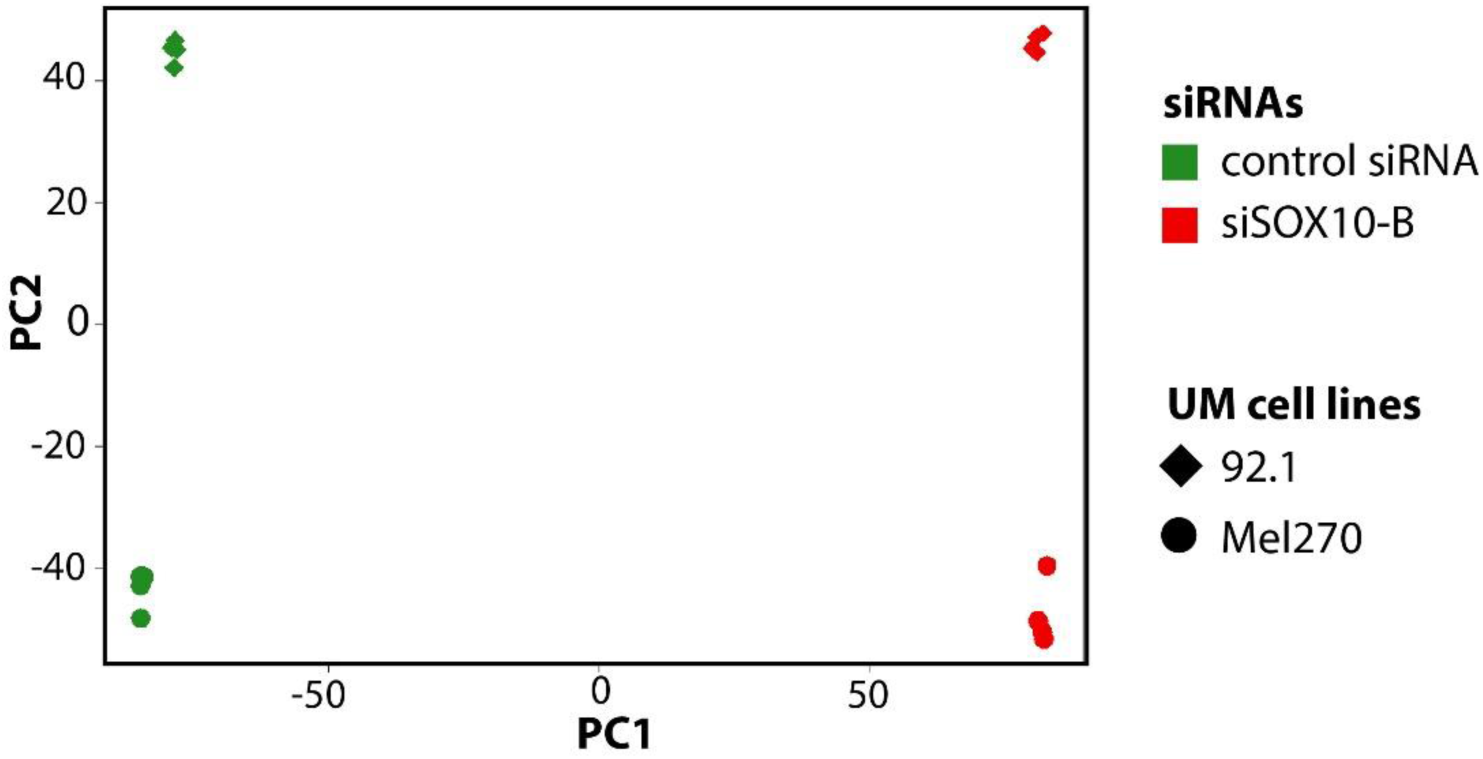
SOX10 knockdown massively alters transcriptomic landscape in UM cells. Two-dimensional principal component (PC) analysis of RNA seq data in UM cell lines 92.1 (diamond) and Mel270 (circle) transfected with control siRNA (green) or siSOX10-B (red) for 24 h, n=4.

**Supplementary Figure S5:**
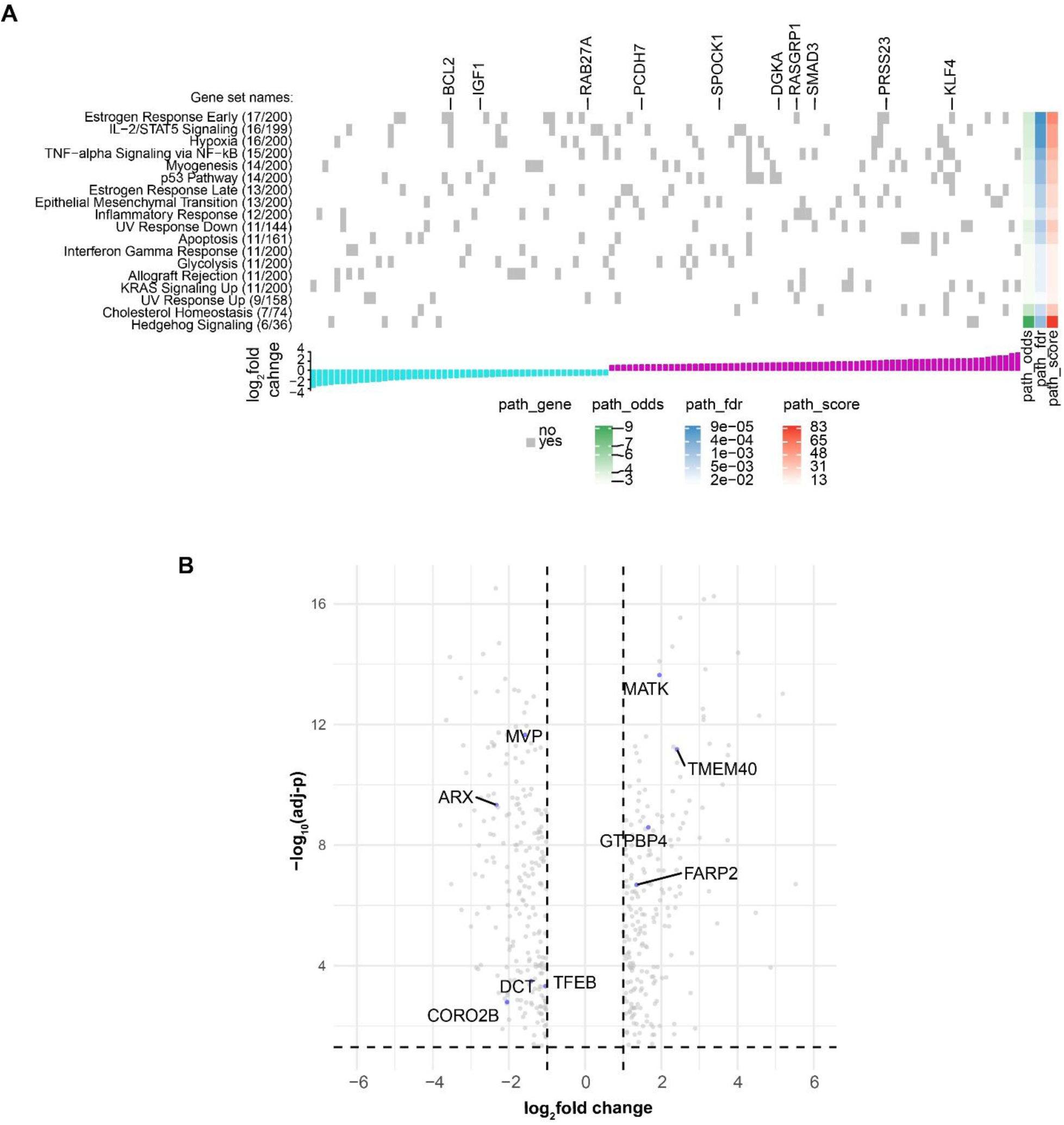
MITF-associated transcriptomic changes after SOX10 knockdown in UM cells. (A) Heat map showing the identified enriched hallmark gene sets from the MSigDB database. Only genes belonging to the identified enriched hallmarks are shown. MITF target genes (n = 2,552) were collected from TRANSFAC and from the literature. All of these genes that were differentially expressed in the SOX10 knockdown condition (n = 435) were used as the input gene list. The gene set names with the number of overlapping genes and the number of total genes in a gene set are shown on the left. Each gray grid in the heat map indicates whether a gene is enriched in a hallmark gene set. The odds ratio (odds), false discovery rate (fdr) and a combined score of each gene set are shown on the right. The bar graph at the bottom shows the log_2_fold change (cyan: downregulated; purple: upregulated) of the genes enriched in the gene sets. The 11 putative targets of MITF are shown at the top. (B) Volcano plot showing the identified differentially expressed MITF target genes after SOX10 knockdown (|log2fold change| ≥ 1 and adj-p ≤ 0.05). Putative MITF target genes that are associated to apoptosis GO terms were labeled and highlighted in blue.

**Supplementary Figure S6:**
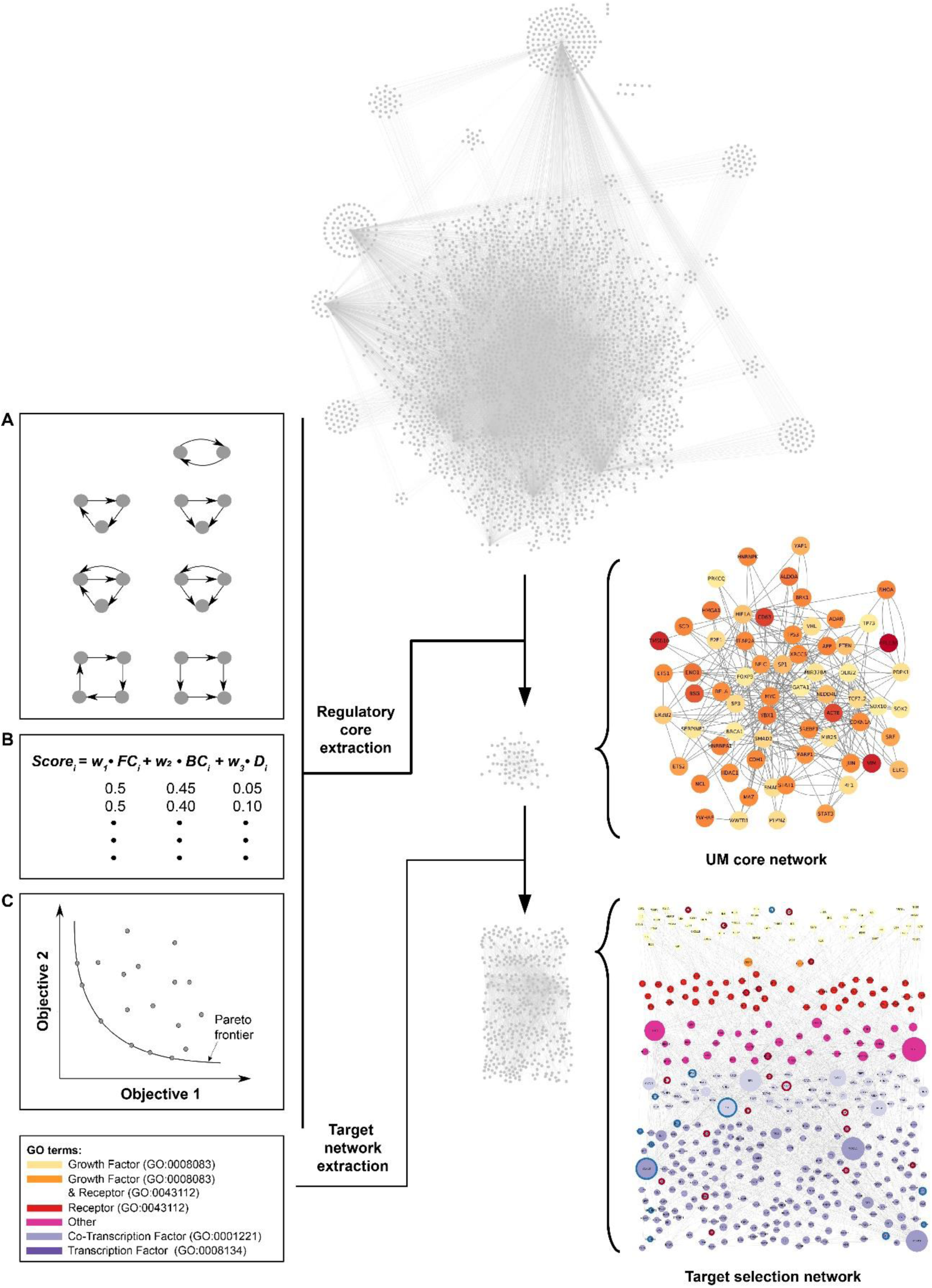
Creating a SOX10-adapted UM protein interaction network to discover new candidates for targeted therapy. Scheme describing the workflow of target selection network development. A SOX10-adapted UM protein interaction network was created by implementing most important intracellular signaling pathways in UM, previously published SOX10 interaction partners, automatically extended with database knowledge using the miRNexpander tool, and expanded by merging with a previously established SOX10 interaction network ^1^. Duplicated nodes and self-loops were removed, the network was pruned with UM gene expression data from public sources ^24^, and nodes with average expression values > 1 were selected. **(A)** Network motifs were explored, **(B)** a weighted ranking score calculated to identify of the most important nodes and interactions, and **(C)** Pareto optimization of the motif scores was used to extract the regulatory core of the network (see Figure S6 for a detailed version). These core genes and their respective first neighbors were extracted from the comprehensive pruned UM network and the nodes were re-annotated according to their basic functionality and reformatted using the CerebralWeb implementation to create the target selection network (see Figure 6 for a detailed version). Additionally, the differential expression values collected from the SOX10 knockdown experiments were projected into the network’s visualization.

**Supplementary Figure S7:**
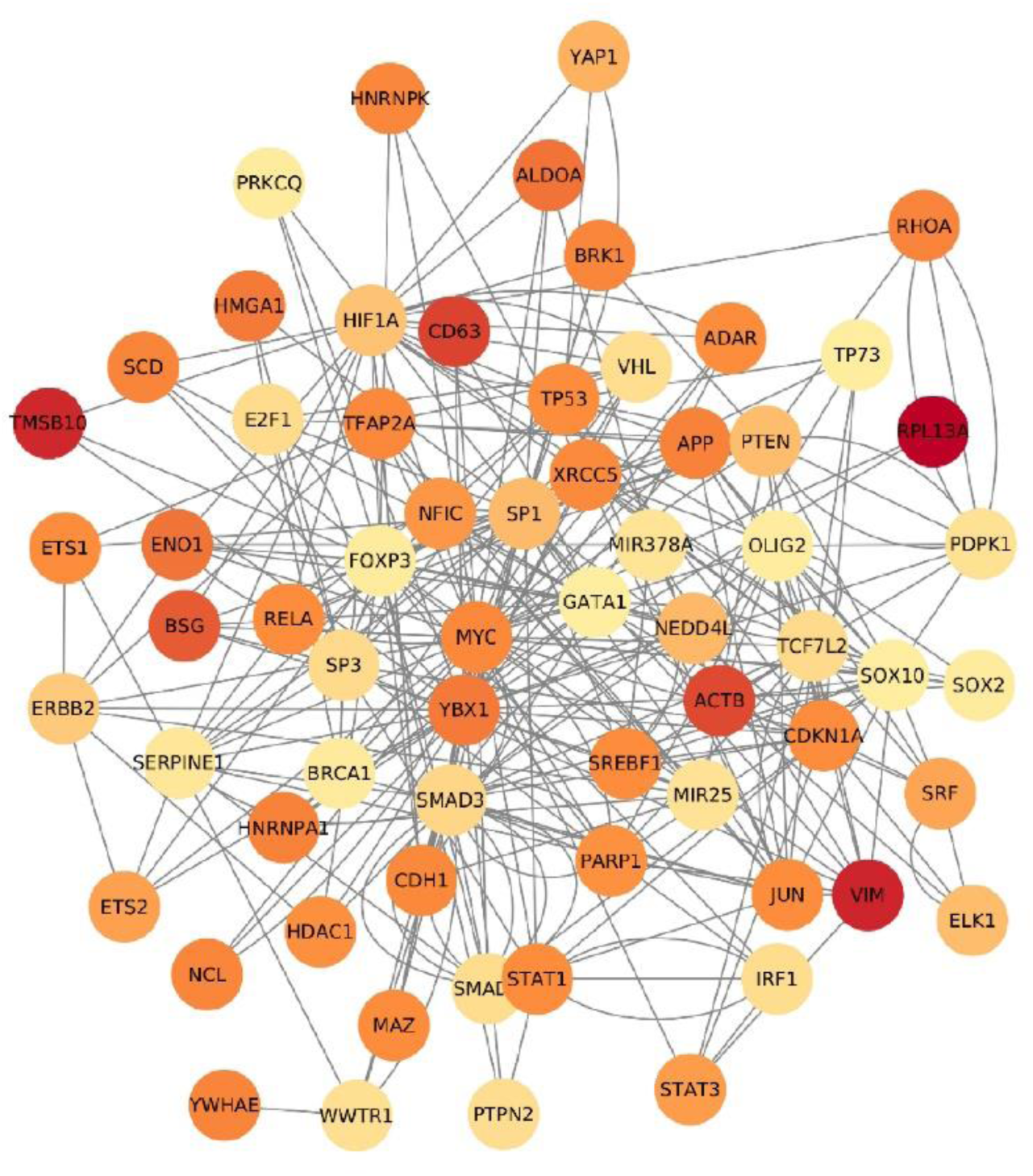
UM core protein interaction network. UM core protein interaction network created by implementing proteins of the most important signaling pathways in UM, known SOX10-protein interactions, and miRNA-mRNA interactions. The 100 top highest-ranked motives comprising 65 nodes and 306 edges are shown.

**Supplementary Figure S8:**
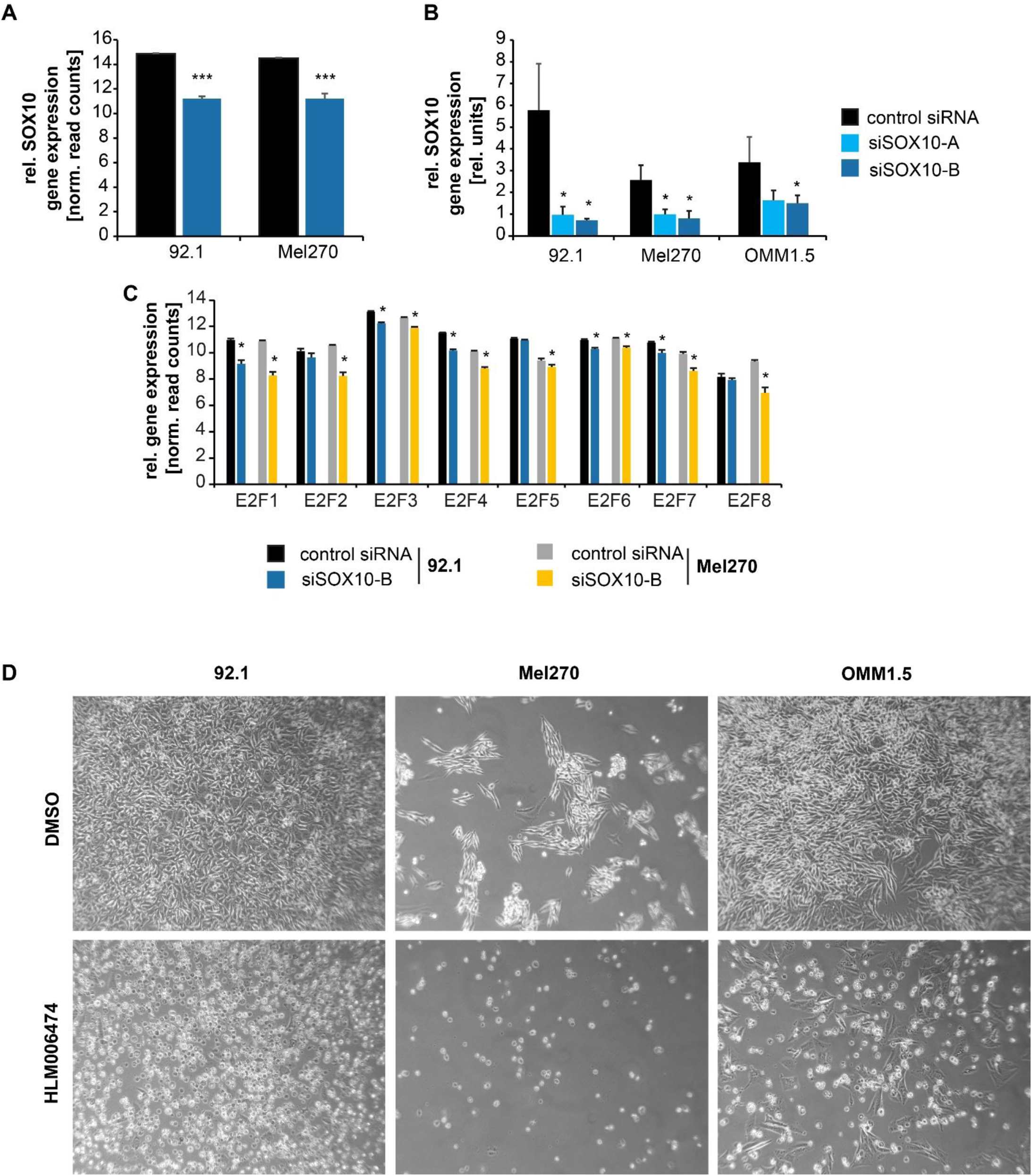
E2F1 as potential candidate for targeted therapy in UM. **(A)** Relative SOX10 gene expression in 92.1 and Mel270 transfected with control siRNA or siSOX10-B for 24 h, determined by RNA seq, normalized read counts, mean ± SD, n = 4. ***: p < 0.001 vs. control siRNA. **(B)** Relative SOX10 mRNA expression levels in 92.1, Mel270, and OMM1.5, normalized to GAPDH expression. Relative expression level of 1205Lu was set to 1, data represents mean ± SD, n=3. *: p < 0.05 vs. control siRNA. **(C)** Relative gene expression of E2F transcription factor family members in normalized read counts in UM cell lines 92.1 and Mel270 transfected with control siRNA or siSOX10-B for 24 h, determined by RNA seq, mean ± SD, n = 4. *: p < 0.05 vs. control siRNA. **(D)** Microscopic images of UM cell lines 92.1, Mel270, and OMM1.5 incubated with 50 µM HLM006474 or DMSO (control) for 96 h, 50fold magnification.

**Supplementary Figure S9:**
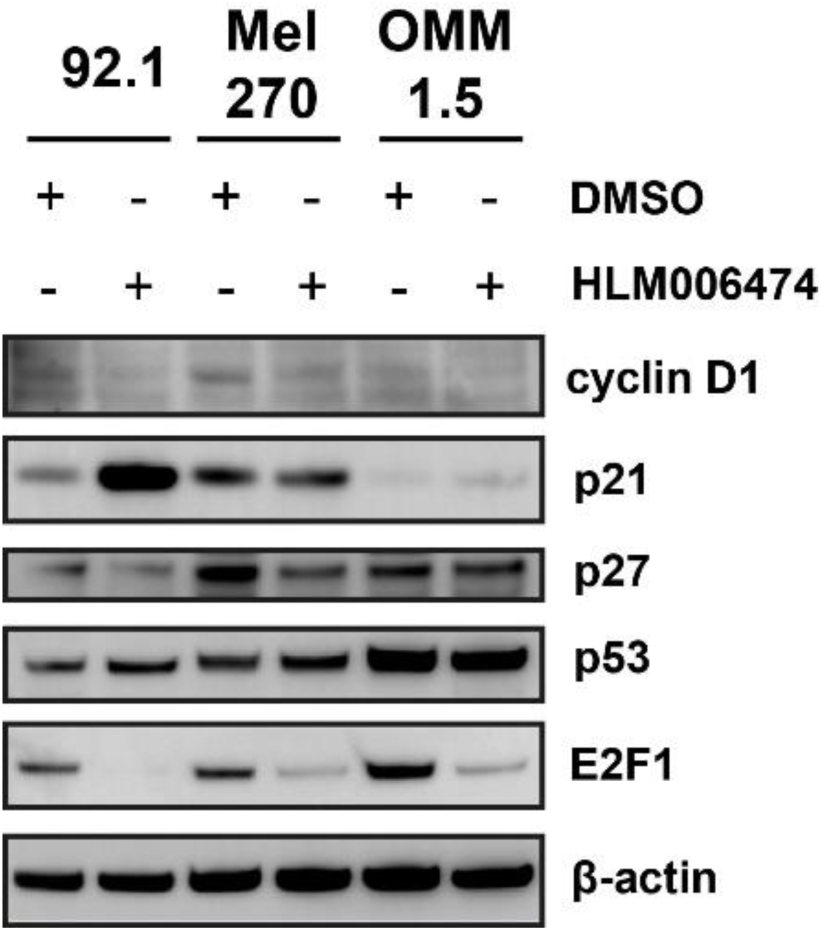
Expression analysis of cell-cycle controlling proteins in UM cells treated with HLM006474. Protein expression of cyclin D1, p21, p27, p53, and E2F1 in 92.1, Mel270, and OMM1.5 incubated with 50 µM HLM006474 or DMSO (control) for 96 h. β-actin served as loading control.

**Supplementary Figure S10:**
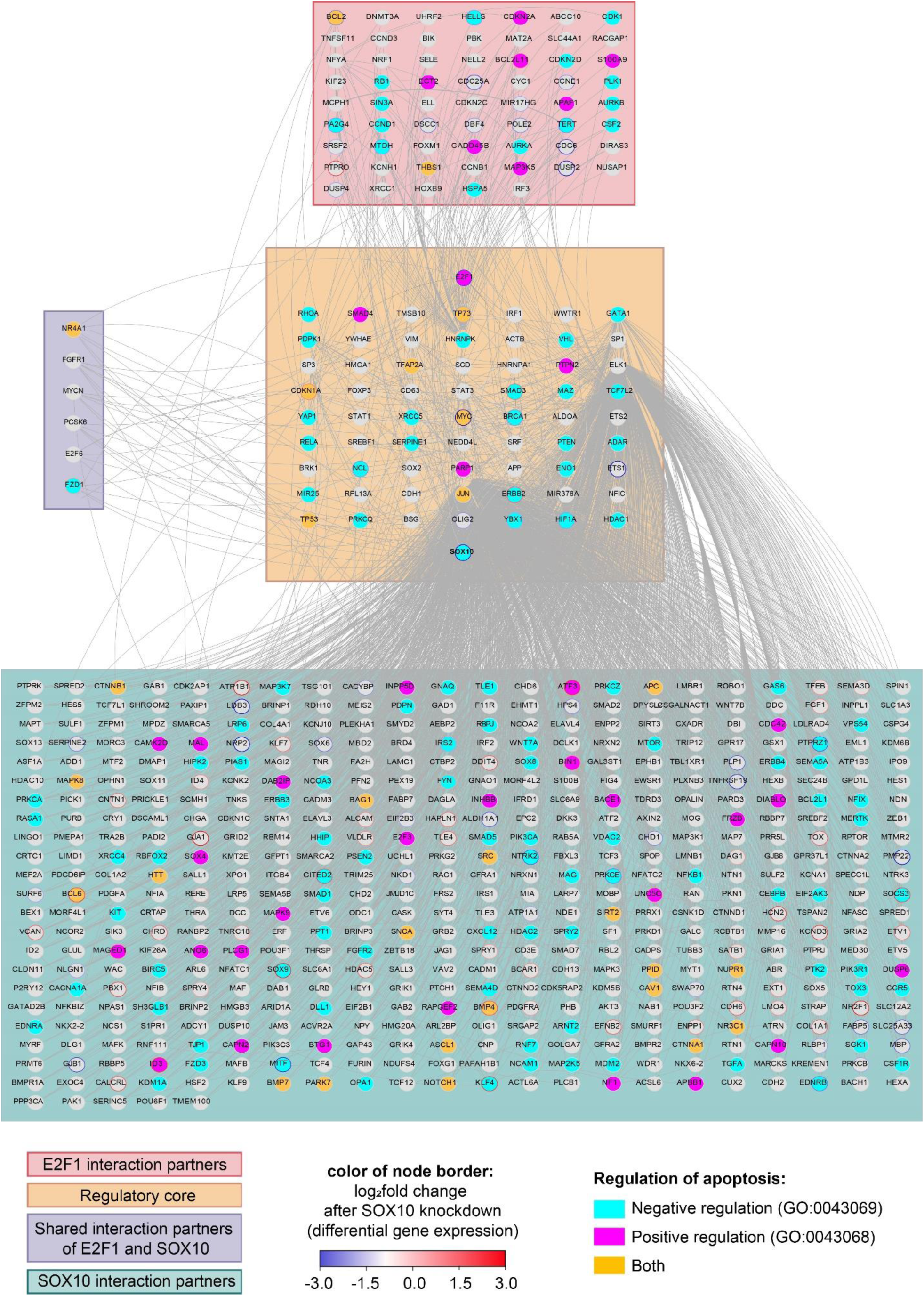
Expanded interaction network for exploring connections between SOX10 and E2F transcription factors. To link SOX10 to the regulation of E2F1 and apoptosis in UM, we use the Cytoscape PlugIn PathLinker ^5^ to obtain from the full UM network all the interaction partners from SOX10 and E2F1. The nodes were colored according to their occurrence in the Gene Ontology annotations for negative regulation of programmed cell death (GO:0043069, cyan), positive regulation of programmed cell death (GO:0043068, pink), or both (orange). The borders of the nodes were colored based on the log2FC expression obtained from the SOX10 K/o experiment. Additionally, we grouped the nodes by either their occurrence in the regulatory core network (light orange), their properties as interaction partners of SOX10 (light green), E2F1 (light red), or both (light purple).

## SUPPLEMENTARY TABLES

**Supplementary Table 1:**
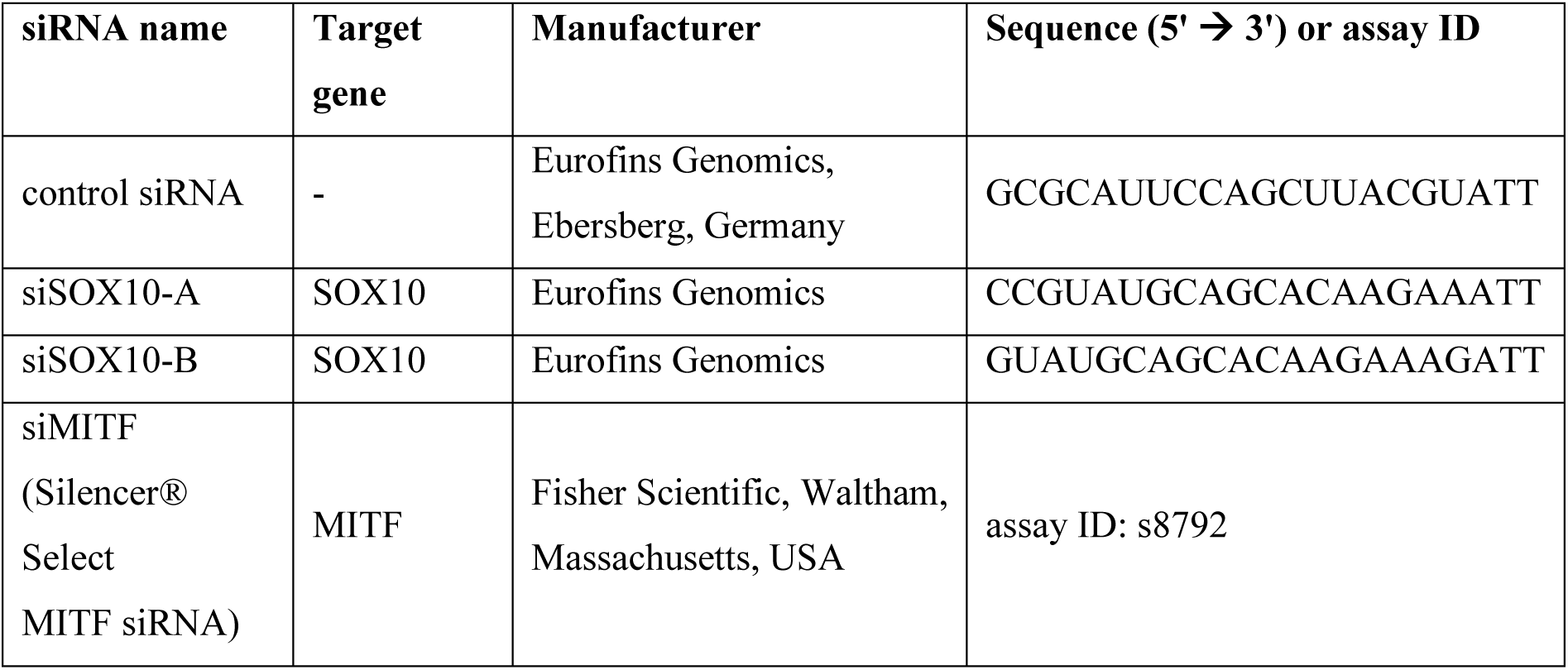
small interfering RNAs.

**Supplementary Table 2:** Intracellular pathways of UM.

| Pathway | REACTOME ID | References |
| --- | --- | --- |
| DNA double-strand break repair | R-HSA-5693532 | 13 |
| Hypoxia response | R-HSA-1234174 | 1,4 |
| Ras-Raf-MEK-ERK-MAPK | R-HSA-5673001, R-HSA-416476 | 1,2,46 |
| Trio-Rho-Rac | R-HSA-9012999 | 6 |
| YAP-Hippo | R-HSA-2028269 | 6 |
| P13K-Akt-mTOR | R-HSA-109704, R-HSA-165159, R-HSA-6807070 | 3,6 |
| PLC-PKC | R-HSA-418217 | 1,4,6 |
| IL-10 | R-HSA-6783783 | 7,8 |
| TGF-beta | R-HSA-170834 | 7,8 |
| IFN-gamma | R-HSA-877300 | 7,8 |
| FasL | R-HSA-75157 | 7,8 |
| PD-1 | R-HSA-389948 | 7,8 |
| MIF | R-HSA-8950732 | 7,8 |
| CD47 | R-HSA-391160 | 7,8 |
| SOX10 | R-HSA-9619665 | 9,10 |

**Supplementary Table 3:** Genes included in the target selection network.

| **HGNC**<br>**gene**<br>**symbol** | **log<sub>2</sub>fold change**<br>**(control siRNA vs. siSOX10-**<br>**B)** | **GO term** |
| --- | --- | --- |
| ABCA2 | 0 | Receptor (GO:0043112) |
| ACHE | 0 | Receptor (GO:0043112) |
| ACTB | 0 | Transcription Factor (GO:0008134) |
| ADAR | 0 | Other |
| ADD1 | 0 | Transcription Factor (GO:0008134) |
| ADM | 0 | Receptor (GO:0043112) |
| AGT | 0 | Growth Factor (GO:0008083) |
| AHR | 0 | Co-Transcription Factor (GO:0001221) |
| ALDOA | 0 | Other |
| ANKRD1 | 0 | Transcription Factor (GO:0008134) |
| ANXA4 | 0 | Transcription Factor (GO:0008134) |
| APBB1 | 0 | Transcription Factor (GO:0008134) |
| APBB3 | 0 | Transcription Factor (GO:0008134) |
| APEX1 | 0 | Transcription Factor (GO:0008134) |
| APOE | 0 | Receptor (GO:0043112) |
| APP | 0 | Other |
| AR | 0 | Co-Transcription Factor (GO:0001221) |
| AREG | 0 | Growth Factor (GO:0008083) |
| ARF1 | 0 | Receptor (GO:0043112) |
| ARID1A | 0 | Transcription Factor (GO:0008134) |
| ARNT | 0 | Transcription Factor (GO:0008134) |
| ARNT2 | 0 | Transcription Factor (GO:0008134) |
| ARNTL | 0 | Transcription Factor (GO:0008134) |
| ASCL1 | 0 | Transcription Factor (GO:0008134) |
| ATF2 | 0 | Transcription Factor (GO:0008134) |
| ATXN2 | 0 | Receptor (GO:0043112) |
| BAIAP2 | 0 | Co-Transcription Factor (GO:0001221) |
| BCL2 | -1.58E+14 | Transcription Factor (GO:0008134) |
| BCL3 | 0 | Transcription Factor (GO:0008134) |
| BCL6 | 2.33873E+13 | Co-Transcription Factor (GO:0001221) |
| BDNF | 0 | Growth Factor (GO:0008083) |
| BEX1 | 0 | Transcription Factor (GO:0008134) |
| BHLHE40 | -1.8383E+14 | Transcription Factor (GO:0008134) |
| BHLHE41 | -2.04804E+14 | Transcription Factor (GO:0008134) |
| BMP4 | 2.05901E+14 | Growth Factor (GO:0008083) |
| BMP7 | 0 | Growth Factor (GO:0008083) |
| BRCA1 | -1.30636E+14 | Other |
| BRK1 | 0 | Other |
| BSG | 0 | Other |
| C1QBP | -1.11834E+13 | Transcription Factor (GO:0008134) |
| CALCRL | 2.08038E+14 | Receptor (GO:0043112) |
| CAV1 | 0 | Receptor (GO:0043112) |
| CBX3 | 0 | Co-Transcription Factor (GO:0001221) |
| CBX5 | 0 | Transcription Factor (GO:0008134) |
| CD36 | 0 | Receptor (GO:0043112) |
| CD63 | 0 | Receptor (GO:0043112) |
| CDH1 | 0 | Other |
| CDKN1A | 1.45737E+14 | Other |
| CDKN2A | 0 | Transcription Factor (GO:0008134) |
| CEACAM1 | 0 | Receptor (GO:0043112) |
| CEBPA | 0 | Transcription Factor (GO:0008134) |
| CEBPB | 0 | Transcription Factor (GO:0008134) |
| CGGBP1 | 0 | Transcription Factor (GO:0008134) |
| CHD6 | 0 | Co-Transcription Factor (GO:0001221) |
| CIITA | -1.59248E+14 | Transcription Factor (GO:0008134) |
| CITED2 | 2.11298E+14 | Transcription Factor (GO:0008134) |
| CREB1 | 0 | Transcription Factor (GO:0008134) |
| CREBBP | 0 | Co-Transcription Factor (GO:0001221) |
| CREM | 0 | Transcription Factor (GO:0008134) |
| CRKL | 0 | Transcription Factor (GO:0008134) |
| CRTC1 | 0 | Transcription Factor (GO:0008134) |
| CRY1 | 0 | Transcription Factor (GO:0008134) |
| CSF1 | 0 | Growth Factor (GO:0008083) |
| CSF2 | 0 | Growth Factor (GO:0008083) |
| CSH1 | 0 | Growth Factor (GO:0008083) |
| CSNK2B | 0 | Transcription Factor (GO:0008134) |
| CTBP2 | 0 | Co-Transcription Factor (GO:0001221) |
| CTCF | 0 | Co-Transcription Factor (GO:0001221) |
| CTDP1 | 0 | Transcription Factor (GO:0008134) |
| CTNNB1 | 0 | Co-Transcription Factor (GO:0001221) |
| CXCL1 | 0 | Growth Factor (GO:0008083) |
| CXCL12 | 1.22672E+14 | Growth Factor (GO:0008083) |
| CXCL8 | 0 | Receptor (GO:0043112) |
| DAXX | 0 | Transcription Factor (GO:0008134) |
| DCAF13 | 0 | Transcription Factor (GO:0008134) |
| DKK1 | 3.00855E+14 | Growth Factor (GO:0008083) & Receptor (GO:0043112) |
| DLG4 | 1.32626E+14 | Receptor (GO:0043112) |
| DLL1 | 0 | Transcription Factor (GO:0008134) |
| DMAP1 | 0 | Transcription Factor (GO:0008134) |
| DNMT3A | 0 | Transcription Factor (GO:0008134) |
| E2F1 | -2.16741E+14 | Transcription Factor (GO:0008134) |
| ECE1 | 0 | Receptor (GO:0043112) |
| EDF1 | 0 | Transcription Factor (GO:0008134) |
| EFNB2 | 1.95838E+14 | Receptor (GO:0043112) |
| EGR2 | 0 | Transcription Factor (GO:0008134) |
| EHMT1 | 0 | Co-Transcription Factor (GO:0001221) |
| EIF4E | 0 | Transcription Factor (GO:0008134) |
| ELK1 | 0 | Transcription Factor (GO:0008134) |
| ENO1 | 0 | Co-Transcription Factor (GO:0001221) |
| ENPP2 | 0 | Transcription Factor (GO:0008134) |
| EP300 | 0 | Co-Transcription Factor (GO:0001221) |
| EPAS1 | 2.40242E+14 | Co-Transcription Factor (GO:0001221) |
| ERBB2 | 1.83698E+14 | Other |
| EREG | 0 | Growth Factor (GO:0008083) |
| ESR1 | 0 | Co-Transcription Factor (GO:0001221) |
| ETS1 | -2.90977E+14 | Co-Transcription Factor (GO:0001221) |
| ETS2 | 0 | Transcription Factor (GO:0008134) |
| EZH2 | -1.0313E+14 | Co-Transcription Factor (GO:0001221) |
| EZR | 0 | Receptor (GO:0043112) |
| FBL | 0 | Transcription Factor (GO:0008134) |
| FCER1G | 0 | Receptor (GO:0043112) |
| FGF1 | 1.49944E+14 | Growth Factor (GO:0008083) |
| FGF11 | 0 | Growth Factor (GO:0008083) |
| FGF2 | 0 | Growth Factor (GO:0008083) |
| FGF21 | 0 | Growth Factor (GO:0008083) |
| FGF22 | 0 | Growth Factor (GO:0008083) |
| FGF9 | -2.53021E+14 | Growth Factor (GO:0008083) |
| FHL2 | 0 | Transcription Factor (GO:0008134) |
| FLNA | 0 | Transcription Factor (GO:0008134) |
| FLT3 | 0 | Transcription Factor (GO:0008134) |
| FMR1 | 0 | Receptor (GO:0043112) |
| FOS | 0 | Co-Transcription Factor (GO:0001221) |
| FOXP3 | 0 | Transcription Factor (GO:0008134) |
| FURIN | 0 | Receptor (GO:0043112) |
| GATA1 | 0 | Co-Transcription Factor (GO:0001221) |
| GATA2 | 0 | Co-Transcription Factor (GO:0001221) |
| GATA3 | 0 | Co-Transcription Factor (GO:0001221) |
| GATA4 | 0 | Transcription Factor (GO:0008134) |
| GDF15 | 0 | Growth Factor (GO:0008083) |
| GFER | 0 | Growth Factor (GO:0008083) |
| GFI1B | 0 | Transcription Factor (GO:0008134) |
| GH1 | 0 | Growth Factor (GO:0008083) |
| GPI | 0 | Growth Factor (GO:0008083) |
| GRB2 | 0 | Receptor (GO:0043112) |
| GRIA1 | 0 | Receptor (GO:0043112) |
| GSK3B | 0 | Transcription Factor (GO:0008134) |
| GTF2H1 | 0 | Transcription Factor (GO:0008134) |
| HAND1 | 0 | Co-Transcription Factor (GO:0001221) |
| HBEGF | 0 | Growth Factor (GO:0008083) |
| HCFC1 | 0 | Transcription Factor (GO:0008134) |
| HDAC1 | 0 | Co-Transcription Factor (GO:0001221) |
| HDAC11 | 0 | Transcription Factor (GO:0008134) |
| HDAC2 | 0 | Transcription Factor (GO:0008134) |
| HDAC3 | 0 | Co-Transcription Factor (GO:0001221) |
| HDAC4 | -1.6619E+14 | Transcription Factor (GO:0008134) |
| HDAC5 | 1.81634E+14 | Co-Transcription Factor (GO:0001221) |
| HDGF | 0 | Co-Transcription Factor (GO:0001221) |
| HES1 | 0 | Co-Transcription Factor (GO:0001221) |
| HEY1 | 0 | Transcription Factor (GO:0008134) |
| HGF | 0 | Growth Factor (GO:0008083) |
| HIF1A | -1.20562E+14 | Co-Transcription Factor (GO:0001221) |
| HIPK2 | 0 | Transcription Factor (GO:0008134) |
| HMGA1 | 0 | Co-Transcription Factor (GO:0001221) |
| HMGA2 | -1.39031E+14 | Transcription Factor (GO:0008134) |
| HNF4A | 0 | Transcription Factor (GO:0008134) |
| HNRNPA1 | 0 | Other |
| HNRNPK | 0 | Other |
| HOXA7 | 0 | Transcription Factor (GO:0008134) |
| HSF1 | 0 | Transcription Factor (GO:0008134) |
| HSPB1 | 0 | Transcription Factor (GO:0008134) |
| ID2 | 0 | Transcription Factor (GO:0008134) |
| ID3 | 2.2935E+14 | Transcription Factor (GO:0008134) |
| ID4 | 1.4108E+13 | Transcription Factor (GO:0008134) |
| IFI16 | 0 | Transcription Factor (GO:0008134) |
| IGF1 | -1.42366E+14 | Growth Factor (GO:0008083) |
| IGF2 | 0 | Growth Factor (GO:0008083) |
| IL10 | 0 | Growth Factor (GO:0008083) |
| IL11 | 0 | Growth Factor (GO:0008083) |
| IL12A | -1.20874E+14 | Growth Factor (GO:0008083) |
| IL12B | 0 | Growth Factor (GO:0008083) |
| IL2 | 0 | Growth Factor (GO:0008083) |
| IL3 | 0 | Growth Factor (GO:0008083) |
| IL4 | 0 | Growth Factor (GO:0008083) |
| IL5 | 0 | Growth Factor (GO:0008083) |
| IL6 | 0 | Growth Factor (GO:0008083) |
| IL7 | 1.64155E+14 | Growth Factor (GO:0008083) |
| INHA | 0 | Growth Factor (GO:0008083) |
| INHBA | 1.6202E+14 | Growth Factor (GO:0008083) |
| INHBB | 0 | Growth Factor (GO:0008083) |
| INSR | 0 | Receptor (GO:0043112) |
| IRF1 | 0 | Other |
| ISL1 | 0 | Transcription Factor (GO:0008134) |
| ITGB1 | 0 | Receptor (GO:0043112) |
| ITGB3 | 0 | Receptor (GO:0043112) |
| JAG1 | 0 | Growth Factor (GO:0008083) |
| JMJD1C | 0 | Transcription Factor (GO:0008134) |
| JUN | 0 | Transcription Factor (GO:0008134) |
| JUNB | 1.18975E+14 | Transcription Factor (GO:0008134) |
| JUND | 1.12594E+13 | Co-Transcription Factor (GO:0001221) |
| KDM1A | 0 | Transcription Factor (GO:0008134) |
| KDM3A | 0 | Transcription Factor (GO:0008134) |
| KDM4C | 0 | Transcription Factor (GO:0008134) |
| KEAP1 | 0 | Transcription Factor (GO:0008134) |
| KIF16B | 0 | Receptor (GO:0043112) |
| KITLG | 0 | Growth Factor (GO:0008083) |
| KLF4 | 2.41387E+13 | Transcription Factor (GO:0008134) |
| KLF5 | 0 | Transcription Factor (GO:0008134) |
| LATS1 | 0 | Transcription Factor (GO:0008134) |
| LDB1 | 0 | Transcription Factor (GO:0008134) |
| LDLR | 0 | Receptor (GO:0043112) |
| LEF1 | 0 | Co-Transcription Factor (GO:0001221) |
| LHX3 | 0 | Transcription Factor (GO:0008134) |
| LMO2 | 0 | Co-Transcription Factor (GO:0001221) |
| LMO4 | 1.34504E+14 | Transcription Factor (GO:0008134) |
| LRP1 | 1.4916E+14 | Receptor (GO:0043112) |
| MACC1 | 0 | Growth Factor (GO:0008083) |
| MAGI2 | 0 | Receptor (GO:0043112) |
| MAPK14 | 0 | Transcription Factor (GO:0008134) |
| MAZ | 0 | Other |
| MED1 | 0 | Co-Transcription Factor (GO:0001221) |
| MED30 | 0 | Transcription Factor (GO:0008134) |
| MEF2A | 0 | Transcription Factor (GO:0008134) |
| MEF2C | 0 | Transcription Factor (GO:0008134) |
| MEIS2 | 0 | Transcription Factor (GO:0008134) |
| METTL23 | 0 | Transcription Factor (GO:0008134) |
| MIA | 0 | Growth Factor (GO:0008083) |
| MIR25 | 0 | Other |
| MIR378A | 0 | Other |
| MTA1 | 0 | Transcription Factor (GO:0008134) |
| MTA2 | 0 | Transcription Factor (GO:0008134) |
| MTDH | -1.1496E+14 | Transcription Factor (GO:0008134) |
| MTF2 | 0 | Co-Transcription Factor (GO:0001221) |
| MTMR2 | 0 | Receptor (GO:0043112) |
| MTOR | 0 | Transcription Factor (GO:0008134) |
| MYC | -3.49736E+14 | Co-Transcription Factor (GO:0001221) |
| MYOCD | 0 | Transcription Factor (GO:0008134) |
| NAB1 | 0 | Transcription Factor (GO:0008134) |
| NAB2 | 0 | Transcription Factor (GO:0008134) |
| NBN | 0 | Transcription Factor (GO:0008134) |
| NCAPG2 | 0 | Transcription Factor (GO:0008134) |
| NCL | -1.33936E+14 | Transcription Factor (GO:0008134) |
| NCOA2 | 0 | Transcription Factor (GO:0008134) |
| NCOA3 | 0 | Transcription Factor (GO:0008134) |
| NCOA7 | -1.18375E+14 | Transcription Factor (GO:0008134) |
| NCOR2 | 0 | Transcription Factor (GO:0008134) |
| NEDD4L | 0 | Other |
| NEK6 | 0 | Co-Transcription Factor (GO:0001221) |
| NFATC1 | 0 | Co-Transcription Factor (GO:0001221) |
| NFATC2 | 0 | Transcription Factor (GO:0008134) |
| NFATC4 | 1.56362E+14 | Transcription Factor (GO:0008134) |
| NFE2L2 | 0 | Co-Transcription Factor (GO:0001221) |
| NFIA | 0 | Transcription Factor (GO:0008134) |
| NFIC | 0 | Other |
| NFYB | 0 | Transcription Factor (GO:0008134) |
| NKX2-1 | 0 | Transcription Factor (GO:0008134) |
| NKX3-1 | 0 | Transcription Factor (GO:0008134) |
| NLK | 0 | Transcription Factor (GO:0008134) |
| NOLC1 | -1.75424E+14 | Transcription Factor (GO:0008134) |
| NOP58 | -1.18859E+14 | Transcription Factor (GO:0008134) |
| NR0B2 | 0 | Transcription Factor (GO:0008134) |
| NR1D1 | 0 | Co-Transcription Factor (GO:0001221) |
| NR1H4 | 0 | Co-Transcription Factor (GO:0001221) |
| NR1I2 | 0 | Transcription Factor (GO:0008134) |
| NR4A1 | 0 | Transcription Factor (GO:0008134) |
| NR4A3 | -1.54273E+14 | Transcription Factor (GO:0008134) |
| NR5A2 | -2.04379E+14 | Co-Transcription Factor (GO:0001221) |
| OGN | 0 | Growth Factor (GO:0008083) |
| OLIG2 | -1.26281E+14 | Other |
| OPHN1 | 0 | Receptor (GO:0043112) |
| OPTN | 0 | Receptor (GO:0043112) |
| PADI2 | 0 | Transcription Factor (GO:0008134) |
| PARK7 | 0 | Transcription Factor (GO:0008134) |
| PARP1 | 0 | Transcription Factor (GO:0008134) |
| PAX2 | 0 | Transcription Factor (GO:0008134) |
| PBX1 | 2.10083E+14 | Co-Transcription Factor (GO:0001221) |
| PCNA | -1.19736E+14 | Transcription Factor (GO:0008134) |
| PCSK9 | 0 | Receptor (GO:0043112) |
| PDGFA | 0 | Growth Factor (GO:0008083) |
| PDGFB | 0 | Growth Factor (GO:0008083) |
| PDGFC | 0 | Growth Factor (GO:0008083) |
| PDGFD | 0 | Growth Factor (GO:0008083) |
| PDPK1 | 0 | Other |
| PER1 | 0 | Co-Transcription Factor (GO:0001221) |
| PER2 | -1.06825E+14 | Co-Transcription Factor (GO:0001221) |
| PGF | 0 | Growth Factor (GO:0008083) |
| PGR | 0 | Co-Transcription Factor (GO:0001221) |
| PIAS1 | 0 | Transcription Factor (GO:0008134) |
| PICK1 | 0 | Receptor (GO:0043112) |
| PIM1 | 0 | Transcription Factor (GO:0008134) |
| PITX1 | 0 | Transcription Factor (GO:0008134) |
| PKN1 | 0 | Transcription Factor (GO:0008134) |
| PPARA | 0 | Co-Transcription Factor (GO:0001221) |
| PPARD | 0 | Co-Transcription Factor (GO:0001221) |
| PPARG | 0 | Co-Transcription Factor (GO:0001221) |
| PPARGC1<br>A | 0 | Transcription Factor (GO:0008134) |
| PPID | 0 | Transcription Factor (GO:0008134) |
| PRDM16 | 0 | Transcription Factor (GO:0008134) |
| PRKCB | -1.09423E+14 | Transcription Factor (GO:0008134) |
| PRKCQ | 0 | Other |
| PRKDC | 0 | Transcription Factor (GO:0008134) |
| PROX1 | 0 | Transcription Factor (GO:0008134) |
| PRRX1 | 0 | Transcription Factor (GO:0008134) |
| PSEN1 | 0 | Receptor (GO:0043112) |
| PSMD9 | 0 | Transcription Factor (GO:0008134) |
| PTEN | 0 | Other |
| PTMA | 0 | Transcription Factor (GO:0008134) |
| PTPN2 | 0 | Transcription Factor (GO:0008134) |
| PURA | 0 | Transcription Factor (GO:0008134) |
| PURB | 0 | Transcription Factor (GO:0008134) |
| RAB5A | 0 | Receptor (GO:0043112) |
| RAMP3 | 0 | Receptor (GO:0043112) |
| RARA | 0 | Co-Transcription Factor (GO:0001221) |
| RB1 | 0 | Transcription Factor (GO:0008134) |
| RBFOX2 | 0 | Transcription Factor (GO:0008134) |
| RBPJ | 0 | Transcription Factor (GO:0008134) |
| REG1A | 0 | Growth Factor (GO:0008083) |
| RELA | 0 | Co-Transcription Factor (GO:0001221) |
| REST | 0 | Transcription Factor (GO:0008134) |
| RHOA | 0 | Other |
| RORA | 0 | Co-Transcription Factor (GO:0001221) |
| RPL13A | 0 | Other |
| RPTOR | 0 | Transcription Factor (GO:0008134) |
| RUNX2 | 1.73045E+14 | Transcription Factor (GO:0008134) |
| RUNX3 | 0 | Co-Transcription Factor (GO:0001221) |
| RUVBL2 | 0 | Transcription Factor (GO:0008134) |
| RXRA | 0 | Co-Transcription Factor (GO:0001221) |
| SCD | 0 | Other |
| SELE | 0 | Receptor (GO:0043112) |
| SERPINE1 | 0 | Other |
| SH3GLB1 | 0 | Receptor (GO:0043112) |
| SIN3A | 0 | Transcription Factor (GO:0008134) |
| SIRT1 | 0 | Transcription Factor (GO:0008134) |
| SIRT2 | 0 | Transcription Factor (GO:0008134) |
| SMAD2 | 0 | Transcription Factor (GO:0008134) |
| SMAD3 | 1.66356E+14 | Co-Transcription Factor (GO:0001221) |
| SMAD4 | 0 | Co-Transcription Factor (GO:0001221) |
| SMARCA1 | 0 | Transcription Factor (GO:0008134) |
| SMURF1 | 0 | Receptor (GO:0043112) |
| SNCA | 0 | Receptor (GO:0043112) |
| SNX1 | 0 | Receptor (GO:0043112) |
| SOX10 | -3.40386E+14 | Transcription Factor (GO:0008134) |
| SOX2 | 0 | Other |
| SOX8 | 0 | Transcription Factor (GO:0008134) |
| SOX9 | 2.78933E+14 | Transcription Factor (GO:0008134) |
| SP1 | 0 | Co-Transcription Factor (GO:0001221) |
| SP3 | 0 | Transcription Factor (GO:0008134) |
| SPI1 | 0 | Transcription Factor (GO:0008134) |
| SRC | 0 | Transcription Factor (GO:0008134) |
| SREBF1 | 0 | Co-Transcription Factor (GO:0001221) |
| SRF | 0 | Transcription Factor (GO:0008134) |
| STAT1 | 0 | Co-Transcription Factor (GO:0001221) |
| STAT3 | 0 | Transcription Factor (GO:0008134) |
| STAT5B | 0 | Transcription Factor (GO:0008134) |
| STK4 | 0 | Transcription Factor (GO:0008134) |
| SUMO1 | 0 | Transcription Factor (GO:0008134) |
| SYK | 0 | Receptor (GO:0043112) |
| TAF12 | 0 | Transcription Factor (GO:0008134) |
| TAF6 | 0 | Transcription Factor (GO:0008134) |
| TAF7 | 0 | Transcription Factor (GO:0008134) |
| TAL1 | 0 | Transcription Factor (GO:0008134) |
| TBP | 0 | Transcription Factor (GO:0008134) |
| TBX2 | 0 | Transcription Factor (GO:0008134) |
| TCF12 | 0 | Transcription Factor (GO:0008134) |
| TCF3 | 0 | Transcription Factor (GO:0008134) |
| TCF4 | 0 | Transcription Factor (GO:0008134) |
| TCF7L2 | 1.08273E+14 | Transcription Factor (GO:0008134) |
| TERT | -1.47884E+14 | Co-Transcription Factor (GO:0001221) |
| TFAP2A | 0 | Other |
| TFDP2 | 0 | Transcription Factor (GO:0008134) |
| TFF1 | 0 | Growth Factor (GO:0008083) |
| TFRC | -1.49927E+14 | Receptor (GO:0043112) |
| TGFA | 0 | Growth Factor (GO:0008083) |
| TGFB1 | 0 | Growth Factor (GO:0008083) & Receptor (GO:0043112) |
| TGFB2 | 0 | Growth Factor (GO:0008083) |
| TGFB3 | 1.08308E+14 | Growth Factor (GO:0008083) |
| THBS4 | 0 | Growth Factor (GO:0008083) |
| THRA | 0 | Transcription Factor (GO:0008134) |
| THRAP3 | 0 | Transcription Factor (GO:0008134) |
| TIMP1 | 0 | Growth Factor (GO:0008083) |
| TLE1 | 0 | Transcription Factor (GO:0008134) |
| TLE4 | 1.73236E+14 | Transcription Factor (GO:0008134) |
| TMSB10 | 0 | Other |
| TP53 | 0 | Transcription Factor (GO:0008134) |
| TP53BP1 | 0 | Transcription Factor (GO:0008134) |
| TP53BP2 | 0 | Transcription Factor (GO:0008134) |
| TP73 | -1.03713E+14 | Transcription Factor (GO:0008134) |
| TPT1 | 0 | Transcription Factor (GO:0008134) |
| TRIP12 | 0 | Transcription Factor (GO:0008134) |
| TWIST1 | 0 | Co-Transcription Factor (GO:0001221) |
| TYMP | 0 | Growth Factor (GO:0008083) |
| UHRF2 | 0 | Transcription Factor (GO:0008134) |
| USF1 | 0 | Transcription Factor (GO:0008134) |
| USF2 | 0 | Transcription Factor (GO:0008134) |
| VDR | -1.41356E+14 | Transcription Factor (GO:0008134) |
| VEGFA | 0 | Growth Factor (GO:0008083) & Receptor (GO:0043112) |
| VEGFC | 1.11517E+14 | Growth Factor (GO:0008083) |
| VGF | 0 | Growth Factor (GO:0008083) |
| VGLL4 | 1.34979E+14 | Co-Transcription Factor (GO:0001221) |
| VHL | 0 | Transcription Factor (GO:0008134) |
| VIM | 0 | Other |
| WDR54 | 0 | Receptor (GO:0043112) |
| WWP2 | 0 | Transcription Factor (GO:0008134) |
| WWTR1 | 0 | Other |
| XRCC5 | 0 | Other |
| YAP1 | 0 | Transcription Factor (GO:0008134) |
| YBX1 | -1.02039E+14 | Other |
| YWHAE | 0 | Other |
| YWHAH | 0 | Transcription Factor (GO:0008134) |
| YY1 | 0 | Transcription Factor (GO:0008134) |
| ZBTB16 | 0 | Co-Transcription Factor (GO:0001221) |
| ZBTB7A | 0 | Co-Transcription Factor (GO:0001221) |
| ZFPM1 | 0 | Transcription Factor (GO:0008134) |
| ZFPM2 | 0 | Transcription Factor (GO:0008134) |
| ZMYND8 | 0 | Transcription Factor (GO:0008134) |

**Supplementary Table 4:**
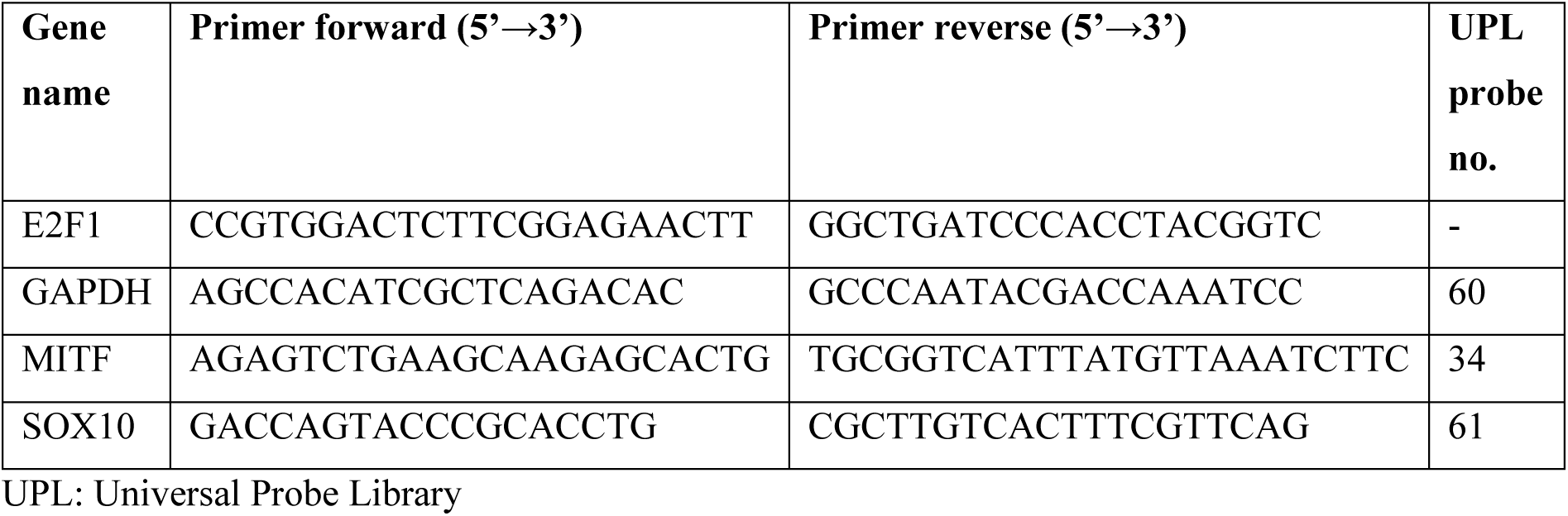
Oligonucleotide primers and hydrolysis probes for qPCR.

**Supplementary Table 5:** Primary antibodies.

| antibody | host | catalog no. | Supplier | Dilution | Research Resource Identifier (RRID) |
| --- | --- | --- | --- | --- | --- |
| anti-β actin | mouse | A1978 | Sigma-Aldrich, Taufkirchen, Germany | 1:5,000 | AB_476692 |
| anti-Akt | rabbit | 9272 | Cell Signaling Technology, Danvers, Massachusetts, USA | 1:1,000 | AB_329827 |
| anti-Bak | rabbit | 3814 | Cell Signaling Technologies | 1:1,000 | AB_2290287 |
| anti-Bax | mouse | sc-20067 | Santa Cruz Biotechnology, Dallas, Texas, USA | 1:1,000 | AB_626726 |
| anti-Bcl-2 | rabbit | 4223 | Cell Signaling Technologies | 1:1,000 | AB_1903909 |
| anti-Bcl-w | rabbit | 2724 | Cell Signaling Technologies | 1:1,000 | AB_10691557 |
| anti-Bcl-xL | rabbit | 2762 | Cell Signaling Technologies | 1:1,000 | AB_10694844 |
| anti-caspase 3 | rabbit | 9662 | Cell Signaling Technologies | 1:1,000 | AB_331439 |
| anti-caspase 9 | rabbit | 9502 | Cell Signaling Technologies | 1:1,000 | AB_2068621 |
| anti-CD71 | rabbit | 13113 | Cell Signaling Technologies | 1:1,000 | AB_2715594 |
| anti-CDK2 | rabbit | 2546 | Cell Signaling Technologies | 1:1,000 | AB_2276129 |
| anti-cyclin A2 | rabbit | 67955 | Cell Signaling Technologies | 1:1,000 | AB_2909603 |
| anti-cyclin D1 | mouse | 33-3500 | Invitrogen, Waltham, Massachusetts, USA | 1:1,000 | AB_86315 |
| anti-cyclin E1 | mouse | 4129 | Cell Signaling Technologies | 1:1,000 | AB_2071200 |
| anti-cytochrome C | mouse | 556433 | BD Pharmingen, Franklin Lakes, New Jersey, USA | 1:500 | AB_396417 |
| anti-E2F1 | mouse | ab4070 | Abcam, Cambridge, United Kingdom | 1:500 | AB_304263 |
| anti-ERK1/2 | rabbit | 9102 | Cell Signaling Technologies | 1:1,000 | AB_330744 |
| anti-FLAG® | rabbit | 14793 | Cell Signaling Technologies | 1:1,000 | AB_2572291 |
| anti-MITF | rabbit | 12590 | Cell Signaling Technologies | 1:1,000 | AB_2616024 |
| anti-Myc-Tag | mouse | 2276 | Cell Signaling Technologies | 1:1,000 | AB_331783 |
| anti-p21 | rabbit | 2947 | Cell Signaling Technologies | 1:1,000 | AB_823586 |
| anti-p27 | mouse | sc-1641 | Santa Cruz | 1:200 | AB_628074 |
| anti-p38 | rabbit | 9212 | Cell Signaling Technologies | 1:1,000 | AB_330713 |
| anti-p53 | mouse | OP43 | Calbiochem/Merck,<br>Darmstadt, Germany | 1:1,000 | AB_564969 |
| anti-PARP | rabbit | 9542 | Cell Signaling<br>Technologies | 1:1,000 | AB_2160739 |
| anti-phospho<br>H2A.X (γH2A.X) | rabbit | 9718 | Cell Signaling<br>Technologies | 1:1,000 | AB_2118009 |
| anti-phospho-Akt<br>Ser473 | rabbit | 9271 | Cell Signaling<br>Technologies | 1:1,000 | AB_329825 |
| anti-phospho-ERK1/2<br>Thr202/Tyr204 | mouse | 9106 | Cell Signaling<br>Technologies | 1:1,000 | AB_331768 |
| anti-phospho-p38<br>MAP Kinase<br>Thr180/Tyr182 | rabbit | 9211 | Cell Signaling<br>Technologies | 1:1,000 | AB_331641 |
| anti-phospho-Rb<br>Ser780 | rabbit | 8180 | Cell Signaling<br>Technologies | 1:1,000 | AB_10950972 |
| anti-phospho-Rb<br>Ser795 | rabbit | 9301 | Cell Signaling<br>Technologies | 1:1,000 | AB_330013 |
| anti-phospho-Rb<br>Ser807/811 | rabbit | 8516 | Cell Signaling<br>Technologies | 1:1,000 | AB_11178658 |
| anti-Rb | rabbit | 9309 | Cell Signaling<br>Technologies | 1:1,000 | AB_823629 |
| anti-SOX10 | mouse | sc-365692 | Santa Cruz | 1:500 | AB_10844002 |

**Supplementary Table 6:**
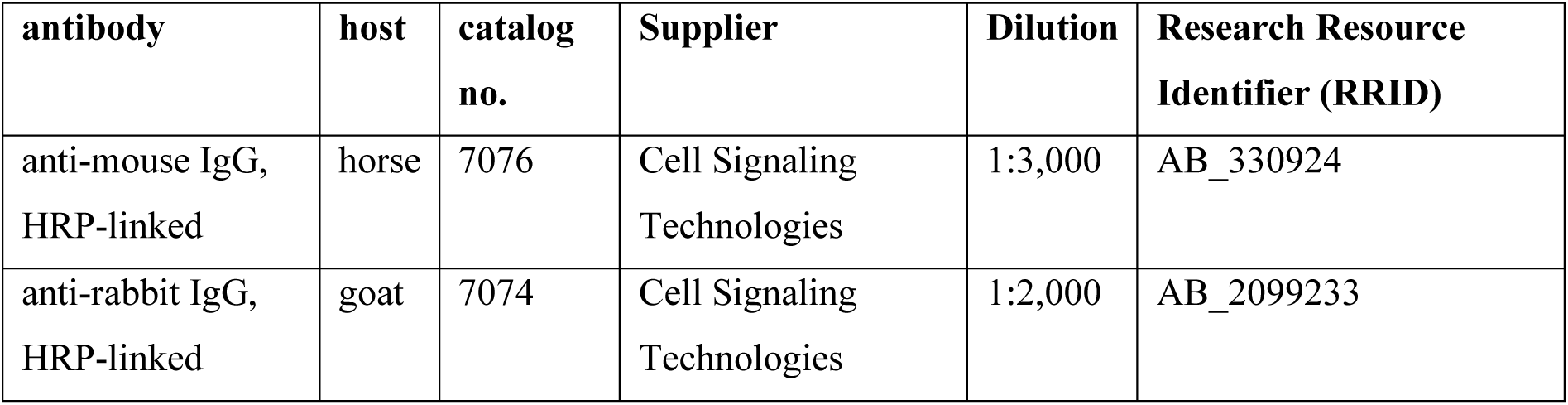
HRP-linked secondary antibodies.

| **antibody** | **host** | **catalog**<br>**no.** | **Supplier** | **Dilution** | **Research Resource**<br>**Identifier (RRID)** |
| --- | --- | --- | --- | --- | --- |
| anti-mouse IgG,<br>HRP-linked | horse | 7076 | Cell Signaling<br>Technologies | 1:3,000 | AB_330924 |
| anti-rabbit IgG,<br>HRP-linked | goat | 7074 | Cell Signaling<br>Technologies | 1:2,000 | AB_2099233 |

**Supplementary Table 7:** References for software, applications, and databases used for network reconstruction.

| **Software, applications or databases**<br>**(version number, if applicable)** | **Reference** |
| --- | --- |
| CellDesigner (version 4.4.2) | 11,12 |
| Minimal Information Required In the Annotation of Models (MIRIAM) | 13 |
| UniProt | 14 |
| Ensembl | 15 |
| miRNexpander | https://github.com/marteber/miRNexpander |
| miRBase | 16 |
| miRTARBase (version 6.1) | 17 |
| miRecords (version 4.5) | 18 |
| HTRIdb (version 1) | 19 |
| TRANSFAC (version 2015.1) | 20 |
| Cytoscape (version 3.8.0) | 21 |
| Gene Expression Omnibus (GEO) | 22 |
| Network Analyzer | 23 |
| NetMatchStar | 24 |

